# Evolution and diversity of biomineralized columnar architecture in early Cambrian phosphatic-shelled brachiopods

**DOI:** 10.1101/2023.06.01.543202

**Authors:** Zhiliang Zhang, Zhifei Zhang, Lars E. Holmer, Timothy P. Topper, Bing Pan, Guoxiang Li

## Abstract

Biologically-controlled mineralization producing organic-inorganic composites (hard skeletons) by metazoan biomineralizers has been an evolutionary innovation since the earliest Cambrian. Among them, linguliform brachiopods are one of the key invertebrates that secrete calcium phosphate minerals to build their shells. One of the most distinct shell structures is the organo-phosphatic cylindrical column exclusive to phosphatic-shelled brachiopods, including both crown and stem groups. However, the complexity, diversity and biomineralization processes of these microscopic columns are far from clear in brachiopod ancestors. Here, exquisitely well-preserved columnar shell ultrastructures are reported for the first time in the earliest eoobolids *Latusobolus xiaoyangbaensis* gen. et sp. nov. and *Eoobolus acutulus* sp. nov from the Cambrian Series 2 Shuijingtuo Formation of South China. The hierarchical shell architectures, epithelial cell moulds, and the shape and size of cylindrical columns are scrutinised in these new species. Their calcium phosphate-based biomineralized shells are mainly composed of stacked sandwich columnar units. The secretion and construction of the stacked sandwich model of columnar architecture, which played a significant role in the evolution of linguliforms, is highly biologically controlled and organic-matrix mediated. Furthermore, a continuous transformation of anatomic features resulting from the growth of diverse columnar shells is revealed between Eoobolidae, Lingulellotretidae and Acrotretida, shedding new light on the evolutionary growth and adaptive innovation of biomineralized columnar architecture among early phosphatic-shelled brachiopods during the Cambrian explosion.

## Introduction

The early Cambrian witnessed a great burst in diversity of animal body plans and biomineralized shell architectures around half a billion years ago (Briggs, 2015; Erwin, 2015, 2020; Budd and Jackson, 2016; Murdock, 2020; Yun et al., 2021; Zhang and Shu, 2021; Zhang et al., 2021c). The novel process of biologically-controlled mineralization producing organic-inorganic composites (hard skeletons), in complex animals has played a vital role in the survival and fitness of early clades (Balthasar and Cusack, 2015; Cuif et al., 2010; Li et al., 2022; Skovsted et al., 2008; Yun et al., 2022), and in turn built the fundamental blocks of complex marine ecosystems (Bicknell and Paterson, 2018; Buatois et al., 2020; Chen et al., 2022; Zhang et al., 2010, 2020). Since the early Cambrian, this adaptive evolution has been demonstrated and continuously preserved in brachiopods, one of the key members of the Cambrian Evolutionary Fauna (Carlson, 2016; Harper et al., 2021, 2017; Sepkoski, 1984). Among them, the phosphatic-shelled brachiopods are some of the most common animals in early Cambrian faunas (Harper et al., 2017). With a high fidelity of preservation and high abundance of biomineralized shells in the fossil record, their morphological disparity, diversity of shell structures and growth patterns, together with ecological complexity are preserved in great detail (Chen et al., 2021; Claybourn et al., 2020; Topper et al., 2018; Zhang, 2018; Z. L. Zhang et al., 2020a, 2021b).

Studying the processes by which organisms form biomineral materials has been a focus at the interface between earth and life sciences (Leadbeater and Riding, 1986; Simonet Roda, 2021). Brachiopods are unique animals in having the ability to secrete two different minerals, calcium phosphate and calcium carbonate, making them the ideal group to further explore the processes of biomineralization (Simonet Roda et al., 2021; Williams, 1977). Hard tissues composed of calcium phosphate with an organic matrix are also largely present in vertebrates, which have remarkably shaped the evolutionary trajectory of life on Earth (Ruben and Bennett, 1987; Wood and Zhuravlev, 2012). The origin of phosphate biomineralization in the evolutionary distant invertebrate brachiopods and vertebrates is still a big mystery in animal evolution (Luo et al., 2015; Neary et al., 2011; Simonet Roda, 2021). Thus, more research is needed in order to understand how to relate hierarchical structure to function in the very early examples of calcium phosphate-based biomineralization processes (Kallaste et al., 2004; Weiner, 2008). It is noteworthy that South China has been considered as one of centres for the origination and early dispersal of phosphatic-shelled brachiopods (Zhang et al., 2021a), and hence, it provides a great opportunity to explore the unique biomineralization process and consequent adaptive evolution of their earliest representatives during the Cambrian radiation.

The shell-forming process of brachiopods, although critical to understanding their poorly resolved phylogeny and early evolution, has long been problematic (Carlson, 2016; Cusack and Williams, 2007; Holmer et al., 2008; Murdock, 2020; Streng et al., 2007; Temereva, 2022; Williams et al., 2004; Williams and Cusack, 1999). The epithelial cells of outer mantle lobes have been considered to be responsible for the shell ornamentation and fabrics of brachiopods (Williams and Cusack, 1999). However, the biologically controlled process of brachiopod shell secretion at the cellular level is still unclear, although organic substrates are observed to be available for biomineral deposition during mantle activity (Williams, 1977). Extensive studies have been conducted on living and fossil shells, but most of them are focused on articulated or carbonate-shelled representatives (Cusack et al., 2010; Griesshaber et al., 2007; Simonet Roda et al., 2021; Simonet Roda, 2021; Ye et al., 2021). By contrast, the linguliform brachiopods with shells composed of an organic matrix and apatite minerals that show extremely intricate architectures and permit exquisite preservation are less studied. The shell structural complexity and diversity, especially of their fossil representatives require further investigation (Butler et al., 2015; Cusack et al., 1999; Streng et al., 2007; Williams and Holmer, 1992; Zhang et al., 2017). The building of shells by microscopic cylindrical columns is a unique feature, which is restricted to the phosphatic-shelled brachiopods and their assumed ancestors (Butler et al., 2015; Holmer et al., 2002, 2008; Holmer, 1989; Skovsted and Holmer, 2003; Williams and Holmer, 2002). This type of columnar shell was previously believed to be exclusively restricted to micromorphic acrotretide brachiopods, a group that demonstrate more complex hierarchical architectures and graded structures compared to simple lamella shell structure in older lingulides (Holmer, 1989; Williams and Holmer, 1992). Further studies however reveal that diverse columnar shell architectures occur in other brachiopod groups, including stem group taxa, such as *Mickwitzia*, *Setatella* and *Micrina* (Butler et al., 2015; Holmer et al., 2008; Skovsted et al., 2010; Skovsted and Holmer, 2003; Williams and Holmer, 2002), the lingulellotretid *Lingulellotreta* (Holmer et al., 2008), the eoobolid *Eoobolus* (Zhang et al., 2021a), and the enigmatic *Bistramia* (Holmer et al., 2008). However, the columnar architectures among the oldest linguliforms and their evolutionary variations have not been studied in detail. Different conditions of fossil preservation with varied taphonomic histories have compounded this issue as different depositional environments, pore-water geochemistry, and subsequent diagenetic and tectonic alteration often obscure the finer details of shell structures (Butler et al., 2015; Holmer et al., 2008; Streng et al., 2007; Ushatinskaya and Korovnikov, 2014; Zhang, 2018).

For the first time, exquisitely well-preserved columnar shell structures are described here from the oldest known eoobolid brachiopods. *Latusobolus xiaoyangbaensis* gen. et sp. nov. and *Eoobolus acutulus* sp. nov. are reported, based on new specimens from the Cambrian Series 2, Shuijingtuo Formation of southern Shaanxi and western Hubei in South China. In this study, the shell architectures, epithelial cell moulds, and the shape and size of cylindrical columns are examined, shedding new light on our understanding of the architecture intricacy, biomineralization process and evolutionary fitness of early phosphatic-shelled brachiopods.

## Results

### Systematic palaeontology

Brachiopoda Duméril, 1806

Linguliformea Williams, Carlson, Brunton, Holmer and Popov, 1996 Lingulata Gorjansky and Popov, 1985

Lingulida Waagen, 1885

Linguloidea, Menke, 1828

Eoobolidae, Holmer, Popov and Wrona, 1996

### Remarks

Holmer et al. (1996, p. 41) established the Eoobolidae to include lingulides characterized by a pitted metamorphic shell and a post-metamorphic shell with pustules. The new taxa described here are assigned to Eoobolidae based on these typical characters. Despite Balthasar’s suggestion to reassign all Eoobolidae members to Zhanatellidae Koneva, 1986, based on the discovery of *Eoobolus* cf. *triparilis* from the Series 2 Mural Formation in the Canadian Rocky Mountains with a pitted metamorphic shell and tuberculate post-metamorphic shell <u>(Balthasar, 2009)</u>, we adhere to Betts’s argument for retaining Eoobolidae <u>(Betts et al., 2019)</u>. Actually, the distinctive features of eoobolids, such as the elevated and divided ventral and dorsal pseudointerareas, are quite different from zhanatellids that are characterized by adpressed dorsal pseudointerareas <u>(Popov and Holmer, 1994; Betts et al.,</u> <u>2019)</u>.

Genus ***Latusobolus*** Zhang, Zhang and Holmer gen. nov.

### Type species

*Latusobolus xiaoyangbaensis* sp. nov., here designated.

### Etymology

From the Latin ‘*latus*’ (wide) with the ending ‘*obolus*’, to indicate the transversely oval outline of both ventral and dorsal valves, morphologically similar to *Obolus*. The gender is masculine.

### Diagnosis

For a full description and discussion of *Latusobolus* gen. nov., refer to Appendix 1.

#### Latusobolus xiaoyangbaensis

Zhang, Zhang and Holmer sp. nov.

Figure 1 ***and*** Appendix 2—figures 1-4, Appendix 3—table 1.

**Figure 1.**
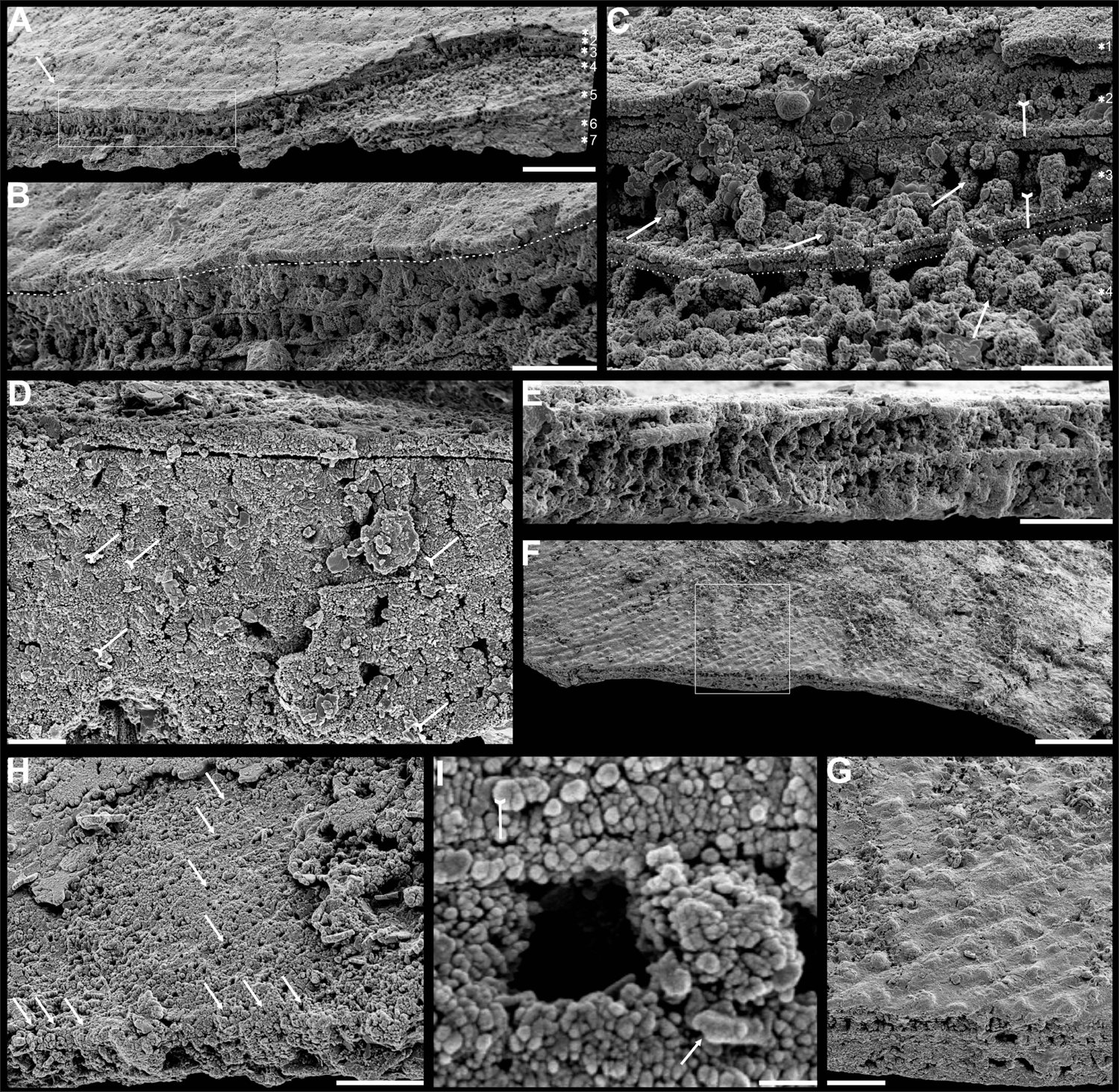
Shell architecture of *Latusobolus xiaoyangbaensis* gen. et sp. nov. from the Cambrian Series 2 Shuijingtuo Formation in southern Shaanxi, South China. **A-C**, ELI-XYB S5-1 BS01. **A**, cross section of a ventral lateral margin, note post-metamorphosis pustules by arrow, primary layer 1 and stacked sandwich columnar units 2-7, box indicates area in **B**. **B**, enlarged view of **A**, showing top primary-underlying secondary layer boundary by dashed line. **C**, enlargement of shell layers 1-4 of **A**, note organic canals of columns (arrows), organic layer (tailed arrows) between two stratiform lamellae by dashed lines. **D**, poorly phosphatised columns of ventral valve, note organic canals by tailed arrows, ELI-XYB S5-1 BR06. **E**, columns of dorsal valve, ELI-XYB S5-1 BS17. **F**, cross section of a ventral lateral margin, show developed post-metamorphosis pustules, box indicates the area in **G**, ELI-XYB S4-2 BO06. **G**, enlarged primary layer pustules and underlying secondary layer columns. **H**, one unit of dorsal stacked columnar architecture with the exfoliation of top primary layer, noting column canals on the stratiform lamella surface by arrows, ELI-XYB S4-2 BO08. **I**, apatite spherules of granule aggregations of ventral columnar shell structure, note granule rods by arrow and thin gap left by the degradation of organic counterparts by tailed arrow, ELI-XYB S4-2 BO06. Scale bars: **A**, 50 µm; **B**, **E**, **G**, 20 µm; **C**, **H**, 10 µm; **D**, 5 µm; **F**, 100 µm; **I**, 1 µm.

#### Etymology

After the occurrence at the Xiaoyangba section in southern Shaanxi, China.

#### Type material

Holotype, ELI-XYB S5-1 BR09 (***Appendix 2—***figure 1M***–P***), ventral valve, and paratype, ELI-XYB S4-2 BO11 (***Appendix 2—***figure 2M***–P***), dorsal valve, from Cambrian Series 2 Shuijingtuo Formation at the Xiaoyangba section (Zhang et al., 2021a) near Xiaoyang Village in Zhenba County, southern Shaanxi Province, China.

**Figure 2.**
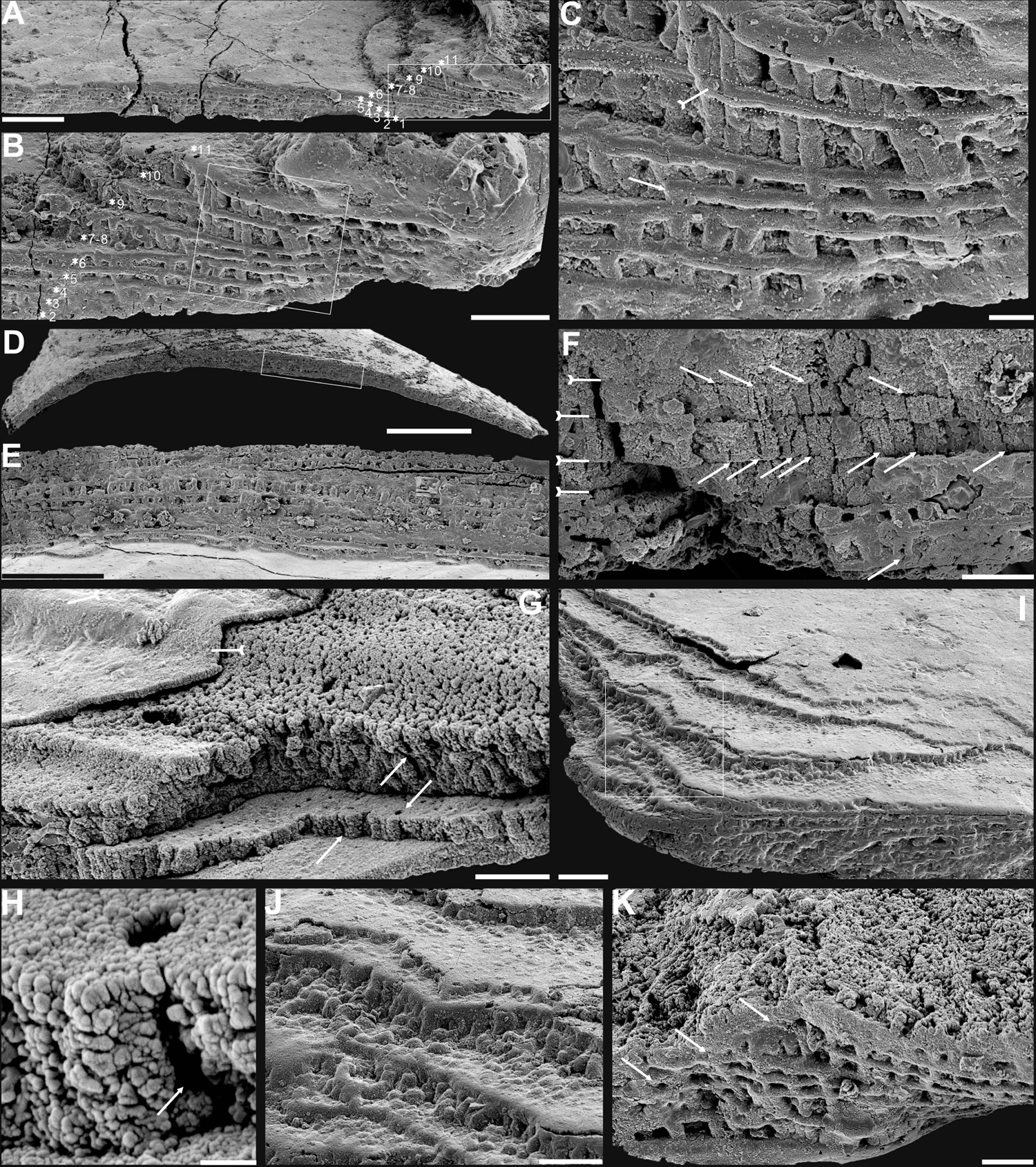
Shell architecture of ventral *Eoobolus acutulus* sp. nov. from the Cambrian Series 2 Shuijingtuo Formation in Three Gorges areas, South China. A-C, ELI-AJH S05 BT12. A, cross section of a ventral lateral margin, note primary layer 1 and stacked sandwich columnar units 2-11, box indicates area in B. B, enlarged view of A. C, enlarged view of B, show thin gap left by the degradation of organic counterparts (tailed arrow) between two stratiform lamellae by dashed lines, the fusion point of two columnar units by arrow. D-F, ELI-AJH 8-2-3 BT02. D, cross section of shell margin, box indicates area in E. E, different preservation condition of columnar architecture. F, poorly phosphatised columns, note the opening of organic canals along the organic membrane by arrows, and space between two stratiform lamellae by tailed arrows. G-H, ELI-AJH 8-2-3 BT03. G, note organic canals on the cross section and surface of stratiform lamella by arrows, and partly exfoliated primary layer by tailed arrow. H, magnified columns composed of granule spherules with canal in G by arrow. I, cross section of shell margin, box indicates area in (J), ELI-AJH 8-2-3 BT04. J, enlarged short columns. K, imbricated columnar architecture (arrows), ELI-AJH S05 BT12. Scale bars: A, E, 50 µm; B, I, 20 µm; C, 5 µm; D, 200 µm; F, G, J, K, 10 µm; H, 1 µm.

#### Diagnosis

As for the genus.

#### Description

For a full description and discussion of *Latusobolus xiaoyangbaensis* gen. et sp. nov., refer to ***Appendix 1***. Genus ***Eoobolus*** Matthew, 1902

#### Type species

*Obolus* (*Eoobolus*) *triparilis* Matthew, 1902 (selected by Rowell, 1965)

#### Diagnosis

See Holmer et al. (p. 41) (Holmer et al., 1996).

***Eoobolus acutulus*** Zhang, Zhang and Holmer sp. nov.

Figure 2 ***and*** Appendix 2—figures 5–7, Appendix 3—table 2.

#### Etymology

From the Latin ‘*acutulus*’ (somewhat pointed), to indicate the slightly acuminate ventral valve with an acute apical angle. The gender is masculine.

#### Type material

Holotype, ELI-AJH S05 BT11 (***Appendix 2—***figure 5E***–H***), ventral valve, and paratype, ELI-AJH S05 1-5-07 (***Appendix 2—***figure 5M), dorsal valve, from Cambrian Series 2 Shuijingtuo Formation at the Aijiahe section (Z. L. Zhang et al., 2016) near Aijiahe Village in Zigui County, north-western Hubei Province, China.

#### Diagnosis

For a full description and discussion of *Eoobolus acutulus* sp. nov., refer to ***Appendix 1***.

### Biomineralized columnar architecture

The shell architectures are exquisitely well-preserved in these newly assigned eoobolid *Latusobolus xiaoyangbaensis* gen. et sp. nov. and *Eoobolus acutulus* sp. nov. Their shell architectures are stratiform in a hierarchical pattern, and consist of primary laminated layer and secondary columnar layer (figures 1 and ***2***). The primary laminated layer is about 1-3 μm thick, composed of compact apatitic lamellae (Figure 1B and **2*G***), while the secondary layer is stratiform with numerous columns disposed orthogonally between a pair of stratiform lamellae (Figures 1B, **C**, **2*B***, ***C***, ***J*** and ***K***; ***Appendix 2—***figure 4I, ***J*** and ***7J***) and looks like being composed of stacked sandwich columnar units. The hollow space in the columns and between lamellae of stacked columnar units may be originally filled with the rich composition of organic material (Figure 1C and D; ***Appendix 2—***figure 4I).

There are 1-3layers of stacked sandwich columnar units developed in *Latusobolus xiaoyangbaensis* gen. et sp. nov. Columns are quite small about 2.4 μm in diameter, ranging from 1.6 μm to 3.4 μm, and about 6 μm in height, ranging from 2.9 μm to 11.9 μm. The central canal in the column ranges from 0.4 μm to 0.9 μm in diameter. The space between the stratiform lamellae is thin, around 0.7 μm, while the stratiform lamellae of columnar units are about 1.4 μm in thickness (***Appendix 3—table 3***).

The maximum number of multi stacked sandwich columnar units increases to 13 in *Eoobolus acutulus* sp. nov. Columns are as small as in *Latusobolus xiaoyangbaensis* gen. et sp. nov., ranging from 1.2 μm to 3.2 μm, and about 4 μm in height. The central canal in the column is small, with the mean diameter of 0.7 μm. The space between stratiform lamellae of stacked columnar units is thin, around 0.6 μm, while the stratiform lamellae are about 1.2 μm in thickness (***Appendix 3—table 3***).

## Discussion

### Diversity of linguliform brachiopod shells

Although the supposed living fossil *Lingula* has long been considered to virtually lack morphological evolutionary changes (Schopf, 1984), more recent studies have shown that lingulide brachiopods have experienced dramatic modifications in many aspects (Liang et al., 2023), including arrangement of internal organs (Zhang et al., 2008), life mode (Topper et al., 2015), shell structure (Cusack et al., 1999), and even genome (Goto et al., 2022; Luo et al., 2015). The complexity and diversity of linguliform shell architecture was increasingly recognised in the pioneering study of Cusack, Williams and Holmer (Cusack et al., 1999; Holmer, 1989; Williams and Cusack, 1999; Williams and Holmer, 1992). Moreover, such complex architectures have a wide distribution in closely related brachiopod groups when they made their first appearance at the beginning of the Cambrian. In connection with an ongoing comprehensive scrutiny of well-preserved linguliform shell ultrastructures from the lower Cambrian limestones of South China, their complexity and diversity hidden in their conservative oval shape is becoming more and more intriguing. However, compared to their ancestral representatives, the shell structure in living lingulides is relatively simple, revealing profound modifications during their long evolutionary history (Cusack et al., 1999; Holmer et al., 2008; Williams, 1977; Williams and Cusack, 1999).

In general, the shells of organo-phosphatic brachiopods are stratiform, composed of an outer periostracum and inner rhythmically-disposed succession of biomineralized lamellae or laminae (Holmer, 1989; Williams, 1977). The organic periostracum, serving as a rheological coat to the underlying shell, is rarely fossilized. However, its wrinkling and vesicular features have largely been preserved as superficial imprints (pits, pustules, fila, grooves, ridges, rugellae, drapes, reticulate networks and spines) on the surface of the primary layer, and are important characters in understanding brachiopod phylogeny (Holmer, 1989; Holmer *et al*., 1996; Cusack *et al*., 1999). By contrast, the structures preserved in the secondary layer have been characterised in fossil and living groups by three ancient fabrics –columnar, baculate, laminated – all of which persist in living shells except for the columnar fabric (Cusack et al., 1999). The tertiary shell layer is well developed in some recent and Palaeozoic lingulides <u>(Holmer, 1989)</u>, but it is not recognised in the early eoobolides.

The primary layer commonly consists of heavily biomineralized compact laminae composed of apatite granules, with a thickness from 2 µm to 20 µm (Williams, 1977) (Figures 1B*, **C*** and ***2G***; ***Appendix 2—***figure 4J). Usually, the concentric growth lines are evenly distributed on the surface of the primary layer of the post-metamorphic shell. By contrast, the surface ornamentation tends to be unevenly distributed, demonstrating a strong phylogenetic differentiation. Superficial pustules are one of the most distinct patterns and are readily recognised in one of the oldest brachiopod groups, the Eoobolidae (Holmer et al., 1996). The pustules are roughly circular in outline, composed of apatite aggregates, and range from 2 µm to 20 µm in diameter (Zhang, 2018) (Figure 1G). Such pustules are also found on early obolid, zhanatellid and acrotheloid shells with relatively wide size variations from 5 µm to 30 µm in diameter (Cusack et al., 1999; Zhang, 2018). Although different in size, the similar pattern may indicate the same secretion regime that originated as vesicles during the very early stages in periostracum secretion (Cusack et al., 1999). The thickness of the underlying secondary layer varies greatly in different brachiopod groups, depending on the shell component and fabric type. Columnar, baculate and laminated fabrics are incorporated into the basic lamination component to form the diverse stratiform successions of the secondary shell layer.

The fossil record reveals that the columnar shell structure is generally preserved in most early linguliforms (Holmer et al., 2008; Streng et al., 2007; Williams, 1977; Zhang, 2018; Z. L. Zhang et al., 2021a, 2020a). It is a multi-stacked sandwich architecture (multi-columnar units are stacked in a vertical direction), which is developed in the earliest linguliform *Eoobolus* with a relatively simple architecture as an early developmental stage (Zhang et al., 2021a).

Each stacked sandwich columnar unit consists of numerous columns disposed orthogonally between a pair of compact stratiform lamellae (Cusack et al., 1999; Holmer, 1989). One to three stacked sandwich columnar units with short orthogonal columns can also be found in *Latusobolus xiaoyangbaensis* gen. et sp. nov. (Figure 1) and *E. incipiens* (Figure 4A), while the number of stacked sandwich units increased in later eoobolids (Figure 2), obolids and lingulellotretids (Figure 4B***–D***). Eventually, the acrotretides developed a more complex columnar architecture with multiple stacked sandwich units (Figure 4F***–J***). Moreover, the columnar shell structures are also found in stem group brachiopods, e.g., *Mickwitzia*, *Setatella* and *Micrina*, but with different column size and numbers of laminae (Butler et al., 2015; Holmer et al., 2002, 2008; Skovsted et al., 2010; Williams and Holmer, 2002). It is assumed here that the columnar architecture may be a plesiomorphic character in linguliform brachiopods, inherited from stem group brachiopods.

### Biomineralization process of organo-phosphatic columnar architecture

Metazoans are known for secreting very different types of biominerals through the process of biological mineralization. This linking of living soft organic tissues with solid earth minerals is a process that has changed the nature of Earth’s fossil archive (Addadi and Weiner, 2014; Lowenstam and Weiner, 1989; Simonet Roda, 2021; Wood and Zhuravlev, 2012). Because of the fine quality of phosphate biomineralization in linguliforms (Cusack et al., 1999; Williams and Cusack, 1999), they can have exquisitely finely preserved shell ultrastructures (figures 3 and ***4***), including epithelium cell moulds (Figure 3C and F***–I***). This permits us to reconstruct the biomineralization process of their apatitic cylindrical columns and address key questions about how these hierarchical structures relate to mechanical functions. Although the biomineralization process of living brachiopods at the cellular level is not well known, biochemical experiments (Cusack et al., 1999, 1992; Lévêque et al., 2004; Williams and Cusack, 1999) have revealed the possibility that the biologically-controlled, organic matrix mediated extracellular mineralization during brachiopod shell secretion. This process can be compared to the hard tissue forming process of mollusc shells and vertebrate teeth (Golub, 2011; Neary et al., 2011; Simonet Roda, 2021).

**Figure 3.**
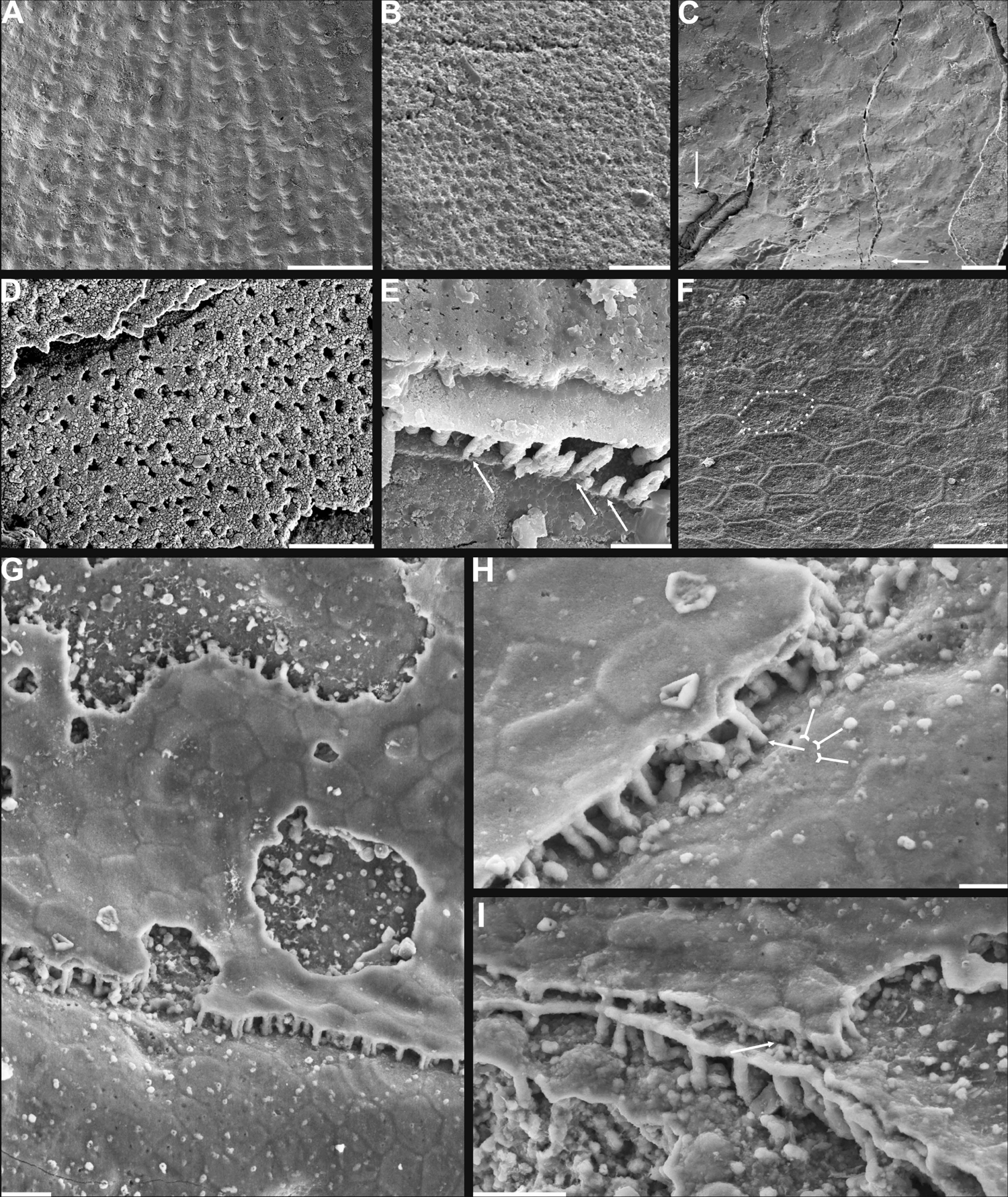
Shell ornamentation, ultrastructure and epithelial cell moulds of Cambrian Series 2 brachiopods. **A**, post-metamorphic pustules of *Latusobolus xiaoyangbaensis* gen. et sp. nov., ELI-XYB S4-3 AU11. **B-D**, *Eoobolus acutulus* sp. nov. **B**, metamorphic hemispherical pits, ELI-AJH 8-2-2 Lin01. **C**, epithelial cell moulds, note column openings on layer surfaces beneath by arrows, ELI-AJH S05 N31. **D**, enlarged column openings on layer surface, ELI-WJP 7 CE05. **E-I**, *Eohadrotreta zhenbaensis*. **E**, partly broken columns, note organic canals by arrows, ELI-AJH F36. **F**, polygonal epithelial cell moulds on valve floor, dash lines indicate margin of one epithelial cell, ELI-WJP 6 R79. **G-I**, ELI-AJH Acro 053. **G**, epithelial cell moulds on dorsal median septum with columns between. **H**, enlarged view of **G**, note rudiment of columns by tailed arrows and one column on epithelial cell margin by arrow. **I**, epithelial cell moulds on stratiform lamella surfaces of successive three stacked sandwich columnar units developed on cardinal muscle areas with columns between (marked by arrow). Scale bars: **A**, **D**, 50 µm; **B**, **E**, **F**, **H**, 10 µm; **C**, **G**, **I**, 20 µm.

Many polygonal structures (Figure 3C and F), preserved on the internal surface of successively alternating laminae (Figure 3G**–*I***), have generally been considered to represent the moulds of epithelial cells (McClean, 1988). The average size of epithelial cells is 20 µm, ranging from 5 to 30 µm in early Cambrian linguliforms (Z. L. Zhang et al., 2016). The rheological vesicle environment may cause the shape variation of epithelia, while different secreting rate could result in different sizes. Generally, smaller epithelial cells had higher secretion rates (Z. L. Zhang et al., 2016). As the active secretion activity of outer epithelial cells, they can be easily embedded in the newly formed inner and outer bounding surfaces of shell laminae (McClean, 1988; Winrow and Sutton, 2012), which left shallow grooves between epithelial margins as intercellular boundaries (Figure 3F). Epithelial cells are preserved as moulds – presumably by the phosphatization of smooth organic sheets (Figure 4G and H) – that had been secreted relatively slowly by the outer plasmalemma of the outer mantle (Cusack et al., 1999).

These active cells are responsible for the secretion of the linguliform periostracum with rheomorphic wrinkling features (Cusack et al., 1999), which left typical microtopography on the external surface of the underlying primary layer (Figure 3A and B). On the other hand, they also secreted linguliform biominerals in a rheological extracellular environment. It is inferred from living *Lingula*, that apatite grows from an amorphous calcium phosphate precursor, which forms the basic crystals of apatite around 5 nm in diameter (Lévêque et al., 2004; Williams and Cusack, 1999). These nanoscale crystals were packaged into apatite granules with an average size of 100 nm (figures 1I and ***2H***; ***Appendix 2—***figure 4K), which acted as the fundamental component building up the hierarchical stratiform shells, including the primary layer and secondary layer (Figure 5). The apatite granules were probably coated or saturated with organic compounds to form granule aggregations or clusters (spherular mosaics) as irregular spherules or rods of about 500 nm (figures 1I and ***2H***; ***Appendix 2—***figures 4J, ***7J*** and ***K***), usually leaving gaps between the aggregation boundaries in fossils after the degradation of organic counterparts (Figure 2H; ***Appendix 2—***figure 4K). These apatite spherules are aggregated in the planar orientation as compact thin lamella less than 4 µm in thickness. Several thin lamellae are closely compacted to form the primary layer (Figure 1B and C; ***Appendix 2—***figure 4I and J). On the other hand, similar nanometre scale networks of spherules are aggregated and organised as orthogonal columns perpendicular to a pair of stratiform lamella surfaces, forming one columnar unit.

Multi-columnar units are stacked in a vertical direction from the exterior to interior to form the secondary layer, applying a stacked sandwich model (Figure 5), which differs from the layer cake model. A very thin gap, commonly less than 1 µm, between each pair of stacked sandwich columnar units is obvious in well preserved specimens (Figures 1C, ***I***, ***2C***, ***F*** and ***4D***; ***Appendix 2—***figures 4I and ***7J***). This is likely indicating an organic membrane acting as an extracellular matrix with functions of a template guiding mineral nucleation (Addadi and Weiner, 2014; Cusack et al., 1999; Lévêque et al., 2004). This is also supported by the almost symmetrical nature of the columnar architecture, revealing a homogeneous organic substrate responsible for the succeeding rhythmic sequence. Although, newly secreted columnar units may succeed the older one unconformably with overlap (Figures 2C, ***3I*** and ***4D***) and be involved in lateral changes of column size (figures 2K and ***4C***), it is supposed to reflect intracellular deviation with the same secretory cycle of the outer mantle as a whole as in living lingulids (Cusack et al., 1999; Williams et al., 1992). The secreting and building processes of the early Cambrian phosphatic-shelled brachiopod columnar shells, secreted by the underlying outer epithelium cells of the mantle lobe, are illustrated in Figure 5.

It is worth noting that, on well preserved specimens nanoscale openings are permeated on the terminal ends of the orthogonal columns (Figures 1C*, **4E*** and ***H***) and surfaces of stratiform lamellae (Figures 1H*, **2G***, ***3D***, ***E***, **4G** and **I**). The openings are rounded with a mean diameter of 600 nm. Observation through natural fractures of shells, show that some canals can be traced continuously through several columnar sequences (Figures 1D, ***2F*** and ***4E***), while in poorly preserved fossils, they are filled with secondarily phosphatised spherules or mosaics (figures 1I and ***2G***), occasionally leaving random gaps (Figures 1D, ***2F*** and ***4I***). These canals are very likely the voids left by degraded organic material, which is confirmed by different taphonomic processes that preserve canals in contemporaneous early Cambrian Burgess Shale–type fossil Lagerstätte in South China (Duan et al., 2021). The regular arrangement of the canal systems and closely related columns reveal that the process is organic matrix-mediated. Furthermore, the even disposition of the organic matrix in columns indicate the existence of rheological and central areas, on which biologically controlled biomineralization took place. It reveals that the nucleation, growth and aggregation of the deposited amorphous calcium phosphate are directed by the same group of epithelial cells (Cusack et al., 1999; Pérez-Huerta et al., 2018; Simonet Roda, 2021;

Weiner and Dove, 2003). In the stacked sandwich columnar units, the empty chambers between each columns (Figure 4F and J) would be originally filled with glycosaminoglycans (GAGs) as in living *Lingula* (Cusack et al., 1999; Williams et al., 1994). These chambers are often filled with coarse spherular mosaics when being secondarily phosphatised and consequently they become indistinguishable from the columns, paired stratiform lamellae and organic membrane (Figures 1D, ***2E***, ***F*** and ***4I***). At the posterior margin of mature shells, especially the ventral pseudointerarea where the vertical component of the growth vector becomes increasingly important in ventral valves, the short columns are succeeded by relatively taller columns, resulting in the change from two stacked sandwich columnar units into one unit during growth anteriorly (Figures 2C, ***K***, ***3I*** and ***4D***). This may demonstrate allometric growth of the shell (Cusack et al., 1999).

The most intriguing and enigmatic phenomenon of skeletal biomineralization is the evolutionary selection of calcium carbonate and calcium phosphate in invertebrates and vertebrates, respectively (Lévêque et al., 2004; Luo et al., 2015). However, the Brachiopoda is a unique phylum that utilises both minerals. The appearance of apatite as a shell biomineral dates back as far as Cambrian Age 2 for stem group brachiopods (Skovsted et al., 2015; Topper et al., 2013; Ushatinskaya, 2002) and persists to the present in living linguliforms (Carlson, 2016). Calcium phosphate can build a relatively less soluble skeletal component compared with calcium carbonate shells, but with the disadvantage of a greater energetic and physiologic cost (Wood and Zhuravlev, 2012). The acquisition of this specific biomineral in phosphatic-shelled brachiopods has been considered an ecological consequence of the globally elevated phosphorous levels during phosphogenic event in the calcite seas with a low Mg:Ca ratio and/or high CO_2_ pressure (Balthasar and Cusack, 2015; Brasier, 1990; Cook and Shergold, 1984; Wood and Zhuravlev, 2012). In such situations, linguliforms were able to utilise the sufficient phosphorous in ambient waters, unlike brachiopod ancestors that possessed an unmineralized shell coated with detrital grains, like *Yuganotheca* found in the Chengjiang Lagerstätte (Zhang et al., 2014). The acquisition of a calcium phosphatic shell may have been an evolutionary response of prey to an escalation of predation pressure during the Cambrian explosion of metazoans (Cook and Shergold, 1984; Wood and Zhuravlev, 2012). Consequently, organic-rich biomineral composites of linguliform brachiopod shells possessed innovative mechanical functions, providing competitive superiority and adaptation on Cambrian soft substrates as well as reducing susceptibility to predation (Wood and Zhuravlev, 2012). Despite the physiological cost of calcium phosphate biomineralization and subsequent reduction in phosphorus levels during post Cambrian period, linguliforms have retained their phosphate shells during dramatical oscillations of seawater chemistry and temperature over 520 million years.

Given the long history of this subphylum, the possession of a phosphatic shell likely has numerous advantages. The innovative columnar architecture can mechanically increase the thickness and strength of the shell by the presence of numerous, stacked thinner laminae, comparable with the laminated fabric seen in obolids (Cusack et al., 1999; Z. F. Zhang et al., 2016). Furthermore, the stacked sandwich columns also increase the strength, flexibility and the ability to resist crack propagation by filling the space between the stratiform lamellae with organic material, comparable with the baculate fabric (Lévêque et al., 2004; Merkel et al., 2009). Thus, the stacked sandwich model of the columnar architecture possesses a greater advantage of mechanical functions and adaptation with a superior combination of strength, durability and flexibility in laminated and baculate fabrics, resembling the colonnaded and reinforced concrete often used in urban construction. New data from nuclear magnetic resonance spectroscopy and X-ray diffraction reveals that apatite in brachiopod shells is highly ordered and thermodynamically stable crystalline and it is more robust in the extremes of moisture, ambient osmotic potential and temperature, unlike the poorly ordered crystal of vertebrate bone (Neary et al., 2011). This type of more efficient and economical shell may also have been responsible for the early diversity of major linguliform brachiopods during the Cambrian explosion, resulting in this group becoming a significant component of the Cambrian Evolutionary Fauna (Bassett et al., 1999; Sepkoski, 1984; Zhang et al., 2008, 2020; Z. L. Zhang et al., 2021b).

### Evolution of stacked sandwich columnar architecture in early brachiopod clades

Evolutionary transformations have repeatedly modified the organo-phosphatic architecture consisting of various aggregates of spherular apatite, held together by a scaffolding of glycosaminoglycan complexes, fibrous proteinaceous struts and chitinous platforms, in linguliform brachiopod shells since the early Cambrian (Cusack et al., 1999). As one of the oldest forms of brachiopod shell architectures, the columnar shell has previously been regarded as an unique character of acrotretide brachiopods (Cusack et al., 1999; Holmer, 1989). However, recent discoveries of columnar shell structures in a diversity of early Cambrian stem group brachiopods have revealed that the same biomineralization strategy is utilised much more widely than previously thought (Butler et al., 2015; Holmer et al., 1996, 2008; Skovsted et al., 2010; Streng et al., 2007; Ushatinskaya and Korovnikov, 2014; Zhang, 2018; Zhang et al., 2021a). This highlights the need for a better understanding of the origin and adaptive modification of stacked-sandwich columnar architectures in early lophophorate evolution.

The Eoobolidae is presently considered to be the oldest known linguliform brachiopods (Zhang et al., 2021a). The biomineralized orthogonal columns in *Eoobolus incipiens* from lower Cambrian Stage 3 probably represents an early and simple shell structure type with a poorly developed secondary columnar layer (Figure 4A). The columns are relatively small with a mean diameter of 1.8 µm ranging from 0.6 to 3.0 µm, and a mean height of 4.1 µm (Figure 6; ***Appendix 3*—*table 3***). The complete secondary layer is only composed of two to three stacked sandwich columnar units, resulting in a shell thickness of about 30 µm. Such a simple shell structure is also developed in the slightly younger *Latusobolus xiaoyangbaensis* gen. et sp. nov. (Figure 1), but with a slightly taller column of about 6.2 µm. From Cambrian Age 4, the number of stacked sandwich columnar unit increases rapidly in the Eoobolidae, growing to as many as 10 stacked sandwich columnar units in *Eoobolus acutulus* sp. nov. (Figure 2), and as many as 10 in *Eoobolus*? aff. *priscus*, and the shell thickness also increases to 50 µm (Streng et al., 2007). However, the size of individual columns keeps within a stable range, around 4 µm in height and 2 µm in diameter. Compared with Eoobolidae, the Lingulellotretidae demonstrates a more developed columnar shell, which has a relatively larger number of stacked sandwich columnar units (up to 20), effectively increasing the shell thickness to 70 µm. The column size is very similar in both *Lingulellotreta malongensis* and *L*. *ergalievi* (Figure 4B and D), and matches that of the contemporaneous *Eoobolus*. But the slightly younger *L*. *ergalievi* has more columnar units, resulting in a thicker shell than *L*. *malongensis*.

**Figure 4.**
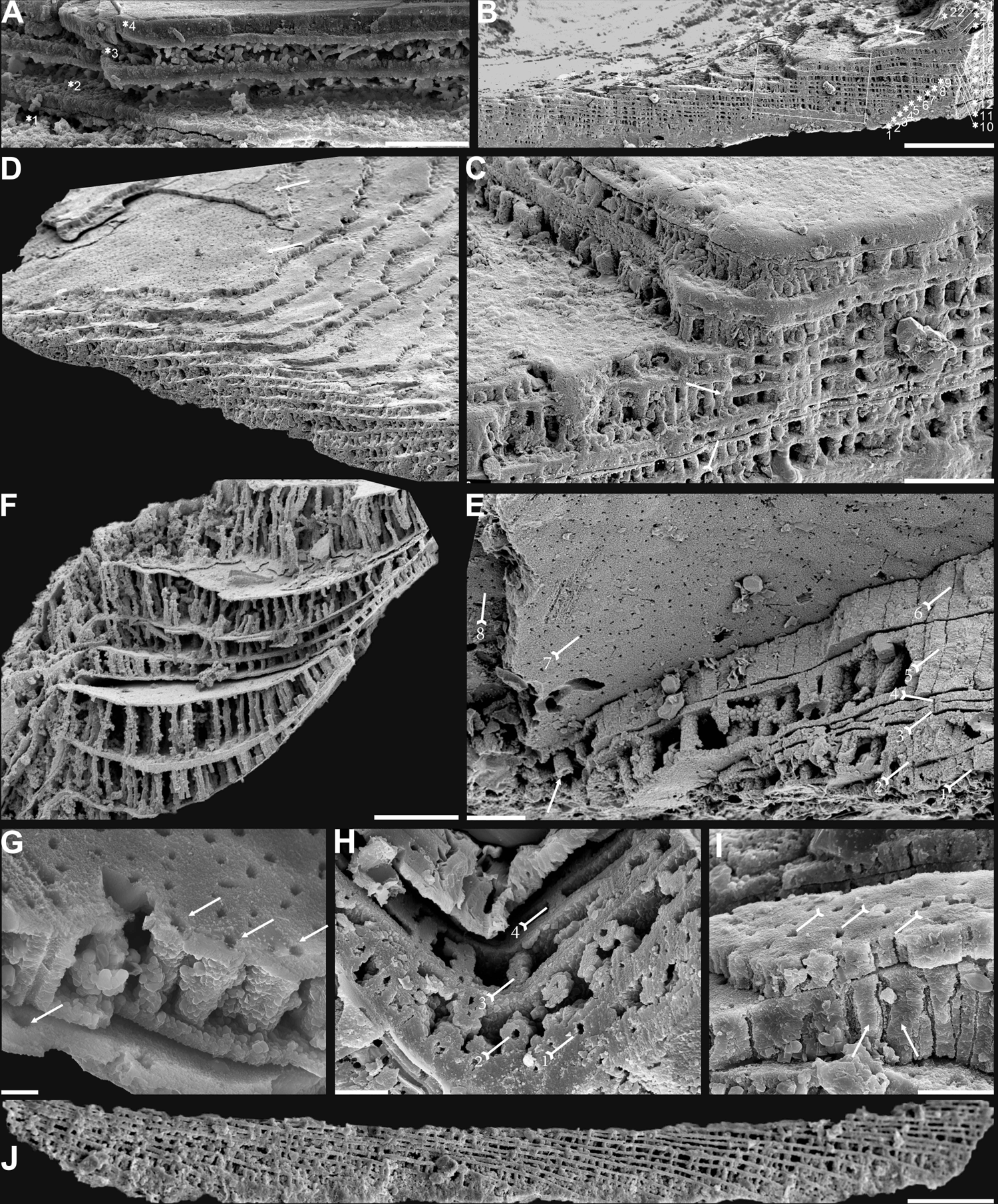
Biomineralized columnar architecture of Cambrian Series 2 brachiopods. **A**, *Eoobolus incipiens*, AJXM-267.5 DT12. **B-D**, *Lingulellotreta ergalievi*, ELI-AJH 8-2-3 CI11. **B**, cross section of shell margin, box indicates area in **C**, note primary layer 1 and stacked sandwich columnar units 2-22, and raised pseudointerarea (tailed arrow). **C**, enlarged view of thin organic layer (tailed arrow) between two stratiform lamellae by dashed lines, the fusion of two stacked columnar units into one by arrow. **D**, imbricated growth pattern of stacked columnar units. **E**, *Palaeotreta zhujiahensis*, note column openings (arrow) on eight successive columnar units by tailed arrows, ELI-AJH 8-2-1 AE09. **F-J**, *Eohadrotreta zhenbaensis*. **F**, relatively taller columns (ca. 20 µm), ELI-AJH 8-2-1 acro16. **G**, apatite spherules of granule aggregations in one columnar unit, note column openings (arrows) on both stratiform lamella surfaces, ELI-AJH S05 E18. **H**, cross section show column openings on 4 successive units by tailed arrows, ELI-WJP 7 AB98. **I**, poorly phosphatised columns (arrows), note openings of organic canals on surface of stratiform lamella by tailed arrows, ELI-AJH S05 I76. **J**, stacked columnar units in an imbricated pattern, ELI-WJP 6 R47. Scale bars: **A**, **E**, **I**, 10 µm; **B**, 100 µm; **C**, **F**, 20 µm; **G**, 2 µm; **H**, 5 µm; **D**, **J**, 50 µm.

**Figure 5.**
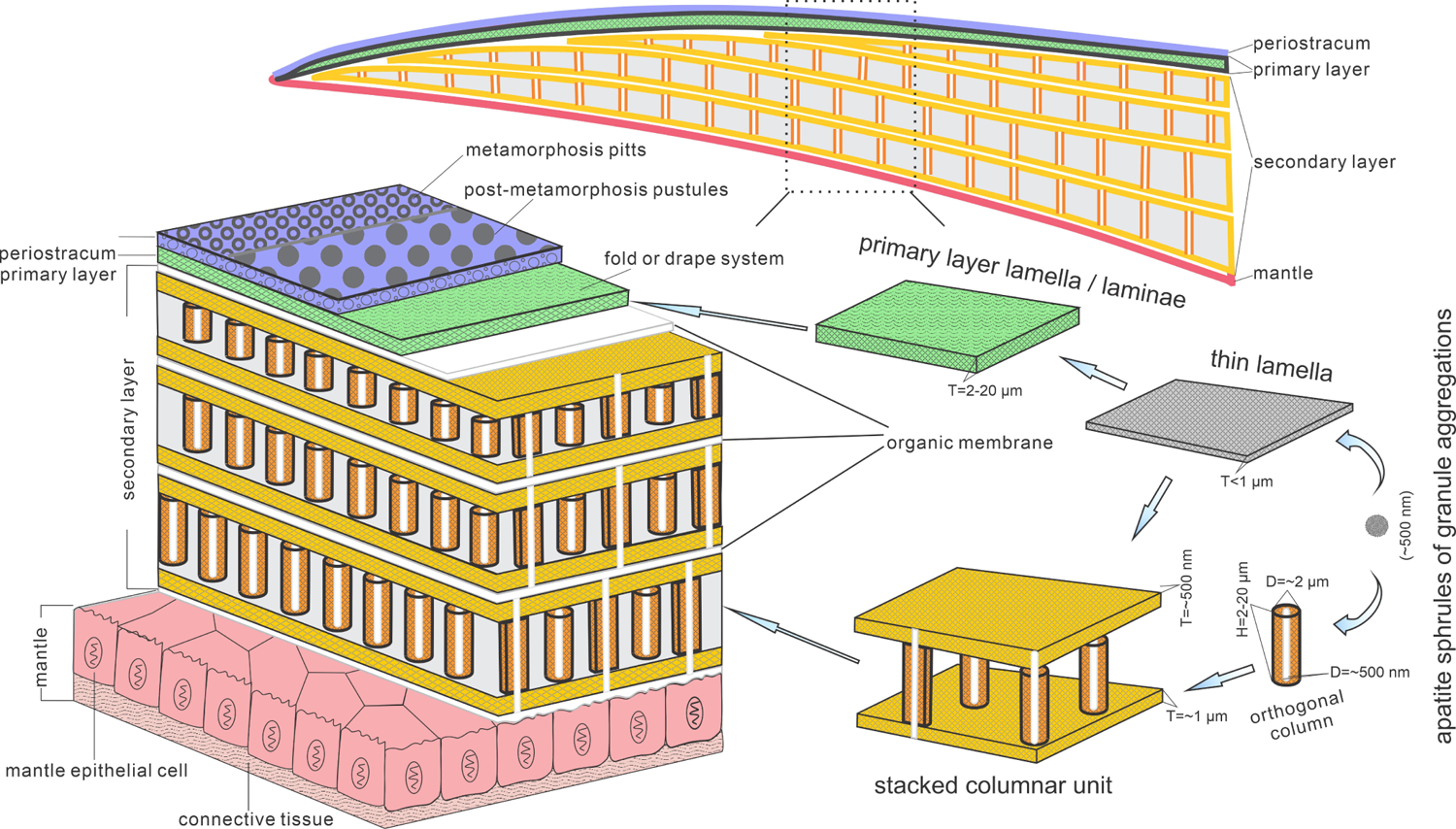
Biomineralization process of typical columnar architecture using the stacked sandwich model of phosphatic-shelled brachiopods. Abbreviation: D=Diameter; H=height; T=Thickness. (modified from (Williams and Holmer, 1992; Z. L. Zhang et al., 2016))

Among all early Cambrian linguliforms with columnar architectures, the acrotretides have developed the most complex shell structure (Figure 4F***–J***). The Cambrian fossil record unveiled a clear pattern of increasing growth (regarding both the diameter and height of the columns and the number of stacked sandwich columnar units) of the columnar architecture in acrotretides: from a very simple type, observed in *Palaeotreta shannanensis* (similar to that of *E. incipiens*) to the slightly more developed structure in *Palaeotreta zhujiahensis* (similar to that of *L. malongensis* (Z. L. Zhang et al., 2020b) to the most advanced architecture observed in *Eohadrotreta zhenbaensis* and younger specimens (Figure 6). The diameter of a single orthogonal column increases about two times in acrotretides compared to eoobolids, whereas the general height of the columns increases to 10 µm in *Eohadrotreta zhenbaensis* and to 29 µm in *Hadrotreta primaeva*, which is about 10 times as high as seen in *Eoobolus variabilis*. Furthermore, the number of columnar units has also increased to about 30, collectively increasing the shell thickness to a maximum value of more than 300 µm in *Eohadrotreta*.

**Figure 6.**
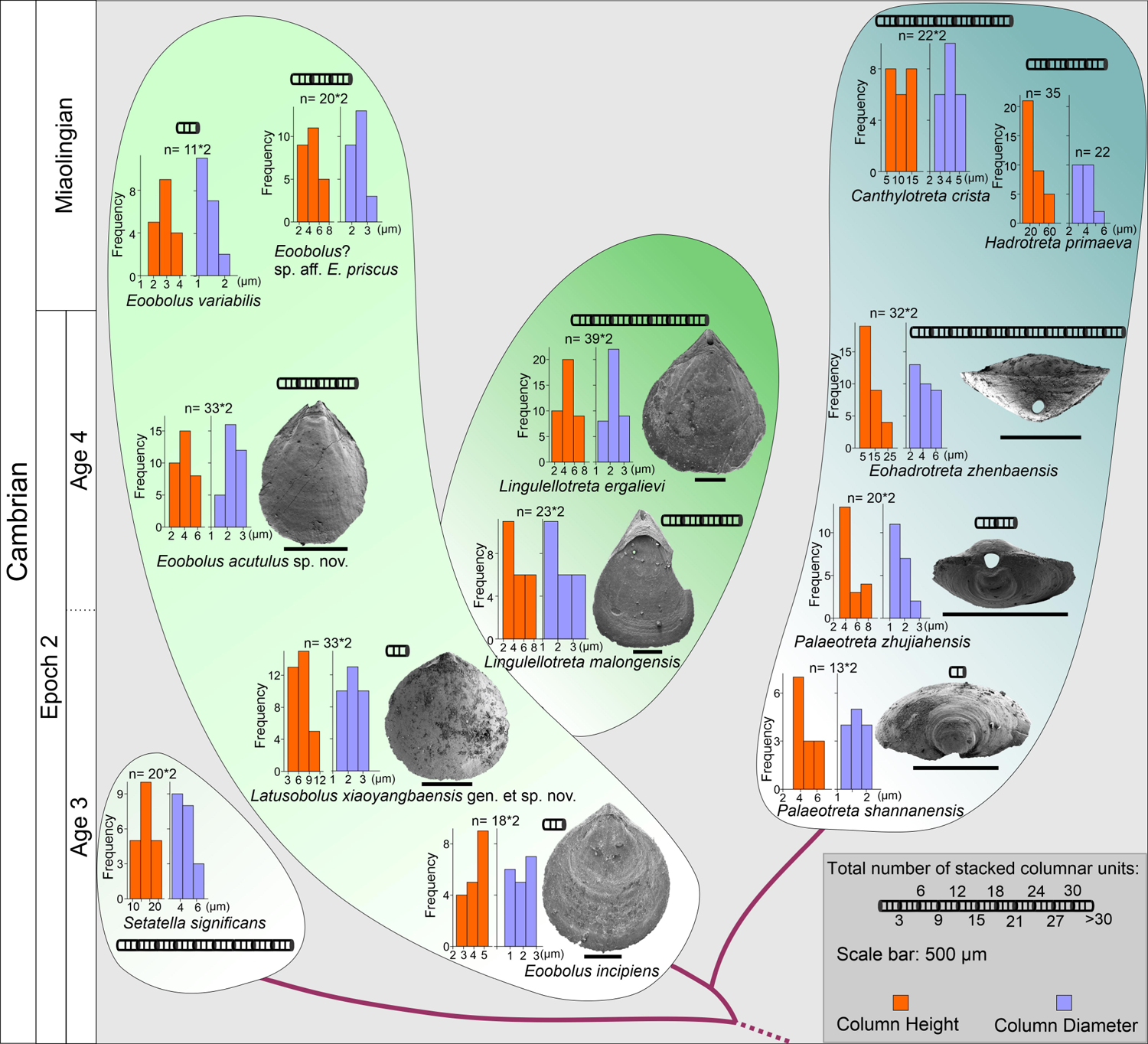
The evolution of stacked sandwich columnar architecture in early Eoobolidae taxa, *Eoobolus incipiens*, *Latusobolus xiaoyangbaensis* gen. et sp. nov., *Eoobolus acutulus* sp. nov., *Eoobolus variabilis*, *Eoobolus*? aff. *priscus*, Lingulellotretidae *Lingulellotreta malongensis*, *Lingulellotreta ergelievi*, Acrotretida *Palaeotreta shannanensis*, *Palaeotreta zhujiahensis*, *Eoohadrotreta zhenbaensis*, *Hadrotreta primaeva*, *Canthylotreta crista* and stem group *Setatella significans*. The height and diameter data of columns are based on data from literature (Skovsted and Holmer, 2003; Streng et al., 2007; Streng and Holmer, 2006; Ushatinskaya and Korovnikov, 2014; Z. L. Zhang et al., 2016, 2020a, 2020b).

The homology of the columnar architecture in early linguliforms outlines a clear picture, most likely representing a continuous transformation between the Lingulida (Eoobolidae and Lingulellotretidae) and Acrotretida. Two groups that represent a major component of early Cambrian benthic communities (Chen et al., 2021; Claybourn et al., 2020; Topper et al., 2015; Zhang et al., 2008; Zhang, 2018; Z. L. Zhang et al., 2020a). The possible occurrence of this shell architecture within the family Obolidae cannot be discounted, as detailed information on the possible columnar shell structures in early Cambrian representatives such as *Kyrshabaktella* and *Experilingula* are poorly known (Cusack et al., 1999; Streng et al., 2007; Williams, 1977). The small size and simple pattern of stacked sandwich columnar architectures remain stable in the Eoobolidae, and this stability likely limits both the shell thickness and overall body size, with ventral and dorsal valves of the family remaining below 6 mm until Miaolingian (middle Cambrian). Stacked sandwich columnar architectures are a character state of the *Lingulellotreta* shell structure as well; however more columnar units are developed that slightly increases the shell thickness and subsequently species of *Lingulellotreta* reach body sizes twice that of Eoobolidae (Zhang et al., 2007). A continuous transformation of anatomic features can be deduced from the evolutionary growth of columnar shells between the two clades. Firstly, the orthogonal columns are markedly developed at the pseudointerarea area of ventral valve of *Lingulellotreta* (Figure 4B), resulting in a greater elevation of the pseudointerarea above the shell floor. It leaves a large amount of space for the posterior extension of the digestive system, which is well protected by the covering mineralized ventral pseudointerarea. This is supported by the discovery of a curved gut under the pseudointerarea of *Lingulellotreta malongensis* in the Chengjiang Lagerstätte (Zhang et al., 2007). Secondly, with the continuous growth of the ventral pseudointerarea, the opening obolide-like pedicle groove is sealed, resulting in a unique pedicle opening exclusively observed in the lingulellotretid brachiopods. Thus, the early pedicle-protruding opening between the ventral and dorsal valves of the Linguloidea is transformed into a new body plan where the pedicle opening is restricted in the ventral valve of lingulellotretids. It supports the scenario that the columnar architecture is monophyletic in at least Linguloidea, and that the slightly younger lingulellotretid columnar architecture was derived from an *Eoobolus-*like ancestor during late Cambrian Age 3.

In another evolutionary direction, acrotretide brachiopods fully utilise the columnar architecture including the derived camerate fabric as shell structure across the whole clade (Streng and Holmer, 2006). The similarity and gradually evolutionary transformations of shell structure from simple forms in lingulides to complex forms in acrotretides suggests the stacked sandwich columnar architecture did not evolve independently in actrotretides. In terms of the derived camerate fabric, more mineralized material is utilised compared to its precursor, the columnar shell structure (Streng et al., 2007). The column size, including height and the number of stacked sandwich columnar units uniformly increase to about 10 times greater in acrotretides than in *E. incipiens and L. xiaoyangbaensis*, since late Cambrian Age 3 (Figure 6). Despite the increase in size of the columns and the number of stacked sandwich columnar units, the whole body of acrotretides is restricted to only millimetre size (Holmer, 1989; Williams et al., 2000). A continuous transformation of anatomic features and shell structure functions can be deduced from the evolutionary growth of columnar shells in early acrotretides. Firstly, the stacked sandwich columns were markedly developed at the pseudointerarea area of the ventral valve, resulting in a greater elevation of the pseudointerarea above the valve floor, compared to that of *Lingulellotreta*.

Secondly, two transformations subsequently changed an obolid-like flat ventral valve (*Palaeotreta shannanensis*) to a cap shape valve (*P. zhujiahensis*) (Z. L. Zhang et al., 2020b), and a conical shape (*Eohadrotreta zhenbaensis*) (Zhang et al., 2018), and eventually to a tubular shape (*Acrotreta*) (Holmer and Popov, 1994). During this evolution, dorsal valves remained relatively flat, showing a limited height profile. The obolid-like ventral pseudointerarea changed from orthocline to catacline and eventually to procline with strongly reduced propareas, while the ventral muscular system moved towards the elevated posterior floor, resulting in the formation of a new apical process (Popov, 1992; Zhang et al., 2018). With the increasing growth of the stacked sandwich columnar shells, the thick organo-phosphatic shell may have increased in strength providing more mechanical support to the conical or tubular valve in a turbulent environment.

Based on the similar evolutionary trajectory, regarding the increasing growth of the shell in Lingulellotretidae and Acrotretida, the formation of their ventral pedicle foramens is very likely homologous, modified from an obolid-like pedicle groove between the two valves. Furthermore, the similarity and continuity in the increasing number and size of the orthogonal columns suggest that columnar architecture is a plesiomorphic character in Linguloidea and Acrotretida. However, the phylogenetic puzzle of whether the columnar architecture is paraphyletic with the baculate fabric in Linguliformea, or even in Lingulata hangs on two pieces of important fossil evidence. The shell structure of the earliest linguliform brachiopods on a global scale needs to be comprehensively investigated based on better preserved fossils. Moreover, more extensive scrutinization of the shell architecture and composition in widely distributed early Cambrian stem group brachiopods is required to conclusively resolve their phylogenetic relationship with linguliforms. The complex shell in stem group taxa *Setatella* and *Mickwitzia*, that are younger and have more advanced columnar shell features than *Eoobolus incipiens*, might reveal the plesiomorphic state of the columnar architecture in Linguliformea (Butler et al., 2015; Holmer et al., 2008; Skovsted and Holmer, 2003; Williams and Holmer, 2002). The accuracy of this assumption depends on future work and whether the columnar shell structure is preserved in older ancestors and other stem group taxa. In another scenario, baculate, laminated and columnar architectures might have originated independently from an unmineralized ancestor like the agglutinated *Yuganotheca* during the early Cambrian (Cusack et al., 1999; Zhang et al., 2014). Regardless of what scenario is true, the origin of the innovative columnar architecture with a stacked sandwich model has played a significant role in the evolution of linguliform brachiopods. The evolutionary diversity of shell architectures would match the general increase in diversity of phosphatic-shelled brachiopods during the Cambrian radiation.

Among them, the micromorphic acrotretides demonstrate the superb application of the columnar architecture combined with its innovative conical shape and possible exploitation of secondary tiering niches (Topper et al., 2015; Wang et al., 2012; Zhang et al., 2008; Z. L. Zhang et al., 2018, 2021b). The fitness of the diminutive body size of acrotretides is likely a trade-off between the increasing metabolic demand of phosphate biomineralization after the Cambrian phosphogenic event and the increased chance of evolutionary survival and adaptation by producing a high mechanical skeleton for protection in the shallow water environment (Cook and Shergold, 1984; Garbelli et al., 2017; Lévêque et al., 2004; Neary et al., 2011; Simonet Roda, 2021; Wood and Zhuravlev, 2012).Their relatively large surface/volume ratio mechanically requires strong support from the composition of stacked sandwich columnar architecture and possibly a relatively lower density of the shell by organic biomineralized material for the secondary tiering life. Such adaptive innovations may account for the flourish of phosphatic-shelled acrotretides in the latter half of the Cambrian, continuing to the Great Ordovician Biodiversification Event, thriving and playing an important role in marine benthic communities for more than 100 million years.

### Material and Methods

The brachiopod material studied here was collected from the Cambrian Series 2 Xihaoping Member of the Dengying Formation and the Shuijingtuo Formation at the Xiaoyangba section of southern Shaanxi (Zhang et al., 2021a), and the Shuijingtuo Formation at the Aijiahe section and Wangjiaping section of western Hubei (Z. L. Zhang et al., 2016). All specimens are recovered through maceration of limestones by acetic acid (∼10%) in the laboratory, and deposited in the Early Life Institute (ELI), Northwest University, China.

Selected specimens were coated and studied further using Fei Quanta 400-FEG SEM at Northwest University, Zeiss Supra 35 VP field emission at Uppsala University and JEOL JSM 7100F-FESEM at Macquarie University. Measurements of length, width and angle of different parts of *Latusobolus xiaoyangbaensis* gen. et sp. nov. and *Eoobolus acutulus* sp. nov. are performed on SEM images of well-preserved specimens by TpsDig2 v. 2.16.

Measurements of diameter and height of orthogonal columns and thickness of different shell layers were performed on SEM images of available adult specimens from this study and previously published literatures by TpsDig2 v. 2.16. Shell thickness was measured at the posterior region of both ventral and dorsal valves of available adult specimens, where the shell displays maximum thickness. The number of columnar units was also counted in the posterior region of available adult specimens. Raw data is provided in ***Appendix 3*—*tables 1-3***.

## Supporting information

Appendix 1-Systematic Palaeontology

## Acknowledgements

We would like to thank Profs. L.E. Popov, G.A. Brock, Y. Cai and C.Y. Cai for insightful discussion, and Q.C. Feng, J.P. Zhai and C.M. Han for sample preparation. Thanks to S. Lindsay and C. Shen at Microscopy Unit at Macquarie University, M. Streng at Uppsala University and Y.L. Pang at Northwest University for assistance with SEM imaging. Thanks also go to two anonymous reviewers and Profs. George Perry and Min Zhu for constructive comments, which greatly improve the manuscript.

## Additional information

### Authors contributions

Zhiliang Zhang, Conceptualization, Investigation, Methodology, Resources, Visualization, Validation, Funding acquisition, Writing – original draft, Writing – review and editing; Zhifei Zhang, Investigation, Resources, Funding acquisition, Writing – review and editing; Lars Holmer, Investigation, Writing – review and editing; Timothy Topper, Investigation, Writing – review and editing; Bing Pan, Writing – review and editing; Guoxiang Li, Investigation, Writing – review and editing.

## Additional files

### Supplementary files

Supplementary file 1. Appendix 1-Systematic Palaeontology.

Supplementary file 2. Appendix 2-figures 1-7.

Supplementary file 3. Appendix 3-table 1.

Supplementary file 4. Appendix 3-table 2.

Supplementary file 5. Appendix 3-table 3.

### Data availability

All data are available in the main text and the supplementary materials.

