## Appendix 1-Systematic Palaeontology for "Evolution and diversity of biomineralized columnar architecture in early Cambrian phosphatic-shelled brachiopods"

### Appendix 1 for:

### Evolution and diversity of biomineralized columnar architecture in the early Cambrian phosphatic-shelled brachiopods

Zhiliang Zhang*, Zhifei Zhang, Lars E. Holmer, Timothy P. Topper, Bing Pan, Guoxiang Li

### Systematic Palaeontology

Phylum Brachiopoda Duméril, 1806

Subphylum Linguliformea Williams, Carlson, Brunton, Holmer and Popov, 1996

Class Lingulata Gorjansky and Popov, 1985

Order Lingulida Waagen, 1885

Superfamily Linguloidea, Menke, 1828

Family Eoobolidae, Holmer, Popov and Wrona, 1996

Genus ***Latusobolus*** Zhang, Zhang and Holmer gen. nov.

**Type species**. *Latusobolus xiaoyangbaensis* sp. nov., here designated.

**Etymology**. From the Latin '*latus*' (wide) with the ending '*obolus*' (Greek coin), to indicate the transversely oval outline of both ventral and dorsal valves, morphologically similar to *Obolus*. The gender is masculine.

**Diagnosis**. Shell transversely suboval or rounded triangular with gently straightened posterior margin, generally 108% wider than long, ventribiconvex. Apical angle is relatively large. Metamorphic shell outlined by a pronounced halo, and ornamented with evenly distributed pitted structures (pits), while post-metamorphic shell ornamented with finely concentric growth lines superposed with round pustules. Columnar shell structure. Ventral pseudointerarea orthocline with shallow and short pedicle groove. Propareas small, with weakly developed flexure lines, slightly raised up. Posterolateral muscle scars and *vascula lateralia* vestigial. Dorsal pseudointerarea orthocline. Propareas small, not raised. Median groove poorly defined, lacking flexure lines. Median tongue short. Umbonal muscle scars weakly impressed. Midian ridge and *vascula lateralia* vestigial.

**Remarks**. The metamorphic pits and post-metamorphic pustules are very unique features for Eoobolidae (Holmer et al., 1996). Although having similar external ornamentation with contemporary *Eoobolus*, the new genus is established for the quite uncommon wide flat shape of both ventral and dorsal valves, while most *Eoobolus* possessing an elongate tongue shape. Compared to *Eoobolus*, the ventral pseudointerarea is smaller and gently raised, posterolateral muscles and pedicle nerve are weakly developed, and the pedicle groove is very short in *Latusobolus*. Furthermore, posterolateral muscles are vestigial, median tongue and median ridge are less developed in dorsal valve of the later. As the columnar architecture is generally discovered in most well studied eoobolid brachiopods (Holmer et al., 2008; Streng et al., 2007; Ushatinskaya and Korovnikov, 2014; Zhang et al., 2021, 2020), it is very likely a new character for family

Eoobolidae. However, the insufficient data of shell architecture in the holotype of *Eoobolus* indicates the comparison with the columnar shells in other eoobolids is not entirely unequivocal. Thus more thorough examinations on shell ultrastructures and ornamentation of early eoobolids and other linguliforms are needed in the future. This will be crucial for better understanding the evolution and phylogeny of brachiopods, and we hope that the comparison study undertaken here can further this aim.

***Latusobolus xiaoyangbaensis*** Zhang, Zhang and Holmer sp. nov.

***Figure 1*** and ***Appendix 2—figures 1-4***, ***Appendix 3—table 1***.

**Etymology**. After the occurrence at the Xiaoyangba section in southern Shaanxi, China.

**Holotype**. ELI-XYB S5-1 BR09 (***Appendix 2—figure 1M–P***), ventral valve.

**Paratype**. ELI-XYB S4-2 BO11 (***Appendix 2—figure 2M–P***), dorsal valve.

**Type locality**. Cambrian Series 2 Shuijingtuo Formation at the Xiaoyangba section (Zhang et al., 2021) near Xiaoyang Village in Zhenba County, southern Shaanxi Province, China.

**Material**. 24 ventral and 17 dorsal valves, ranging from 624 μm to 2325 μm in length and from 669 μm to 2417 μm in width (***Appendix 2—figures 1***, ***2*** and ***Appendix 3—table 1***) from Cambrian Series 2 Shuijingtuo Formation at the Xiaoyangba section, South China.

**Diagnosis**. As for the genus.

**Description**. Shell slightly ventribiconvex, transversely suboval to rounded triangular with gently acuminate posterior margin and rectimarginate anterior commissure, about 108% as wide as long (***Appendix 2—figures 1*** and ***2***). Metamorphic shell ornamented by regularly disposed hemispherical pits, uniform in size of about 0.5 μm, ranging from 0.3 μm to 0.7 μm in diameter (***Appendix 2—figure 3H*** and ***Appendix 3—table 1***). The pronounced halo marks the boundary between metamorphic shell and post-metamorphic shell (***Appendix 2—figure 3A–E***). A narrow belt of about 50 μm width, outside the metamorphic shell lacks pustules and may belong to the neanic shell (***Appendix 2—figure 3A*** and ***D***). Post-metamorphic shell fully covered with fine concentric growth lines superposed by finely pustular ornamentation, (***Appendix 2—figures 1I*** and ***2M***). Round pustules have mean diameter of 6.5 μm, ranging from 2.3 μm to 12.6 μm (***Appendix 2—figure 3F*** and ***G***), packed in concentric rows and rarely in radial pattern (***Appendix 2—figure 3F***).

Ventral valve obtuse with apical angle of 129° on average, rounded triangular, and gently convex in sagittal profile, about length 95% width and depth 18% length, with maximum width slightly anterior to mid-length and maximum height slightly posterior to mid-length (***Appendix 3—***

***table 1***). Pseudointerarea, small, orthocline, occupying 11% valve length and 44% valve width, with a shallow, short, subtriangular pedicle groove of about 112 μm in length and 120 μm in width, with about 64% length of the pseudointerarea (***Appendix 2—figure 4A*** and ***C–E***). Lateral sides of the pedicle groove divergent anteriorly at about 34°. Propareas narrow, raised above the valve floor, divided into two parts by shallow flexure lines, while the internal part of proparea is slightly larger than the external part (***Appendix 2—figures 1J*** and ***4A***). Ventral valve interior with a vestigial visceral area, bisected by the divergent pedicle nerve impression terminated anteriorly at about 36% of valve length (***Appendix 2—figure 4E***). Paired posterolateral muscle scars weakly developed underneath the raised propareas (***Appendix 2—figure 4A*** and ***E***). *Vascula lateralia* weakly developed only on large valves, submarginal, divergent proximally (***Appendix 2—figures 1O*** and ***4A***). Metamorphic shell 223 μm long and 275 μm wide on average, bounded by a pronounced halo (***Appendix 2—figure 3A–C***). Possible protegulum noted as a distinct mound about 50 μm across located posteromedially and bearing faint folds anterolaterally (***Appendix 2—figure 3B***).

Dorsal valve obtuse with apical angle of 136° on average, rounded triangular, and slightly convex in sagittal profile, about 93% shorter than wide and depth 20% length, with maximum width slightly anterior to mid-length and maximum height slightly posterior to mid-length (***Appendix 3—table 1***). Pseudointerarea, small, orthocline, occupying 9% valve length and 41% valve width, with a shallow, short, subtriangular median groove of about 104 μm in length and 292 μm in width, with about 83% length of the pseudointerarea (***Appendix 2—figure 4F*** and ***G***). Median groove divergent anteriorly with an average angle of 112°. Propareas narrow, not raised above the valve floor. Dorsal valve interior with a vestigial visceral area. Median ridge narrow, weakly developed on large valves (***Appendix 2—figure 4G*** and ***H***). Paired posterolateral muscle scars weakly developed anterolaterally to the median groove (***Appendix 2—figures 2N*** and ***4G***). Metamorphic shell 198 μm long and 260 μm wide on average, bounded by a pronounced halo (***Appendix 2—figure 3D*** and ***E***). Possible protegulum noted as a distinct mound about 50 μm across located posteromedially and bearing faint folds anterolaterally (***Appendix 2—figure 3E***).

Shell structure stratiform, consisting of primary laminated layer and secondary columnar layer (***Figure 1***). The primary laminated layer is composed of compact apatitic lamellae, about 3 μm thick (***Figure 1B***), while the secondary layer is composed of stacked sandwich columnar units (totalled 1-3 units), including numerous columns disposed orthogonally between a pair of stratiform lamellae (***Figure 1B*** and **C**; ***Appendix 2—figure 4I*** and ***J***). The hollow space in the columns and between lamellae of stacked columnar units probably indicates the rich composition of organic material (***Figure 1C*** and ***D***; ***Appendix 2—figure 4I***). Columns are quite small about 2.4 μm in diameter, ranging from 1.6 μm to 3.4 μm, and about 6 μm in height, ranging from 2.9 μm to 11.9 μm. The central canal in the column small, ranging from 0.4 μm to 0.9 μm in diameter. The central space between the stratiform lamellae, thin, around 0.7 μm, while the stratiform lamellae of columnar units about 1.4 μm in thickness (***Appendix 3—table 3***).

Genus ***Eoobolus*** Matthew, 1902

Type species. *Obolus* (*Eoobolus*) *triparilis* Matthew, 1902 (selected by Rowell, 1965)

Diagnosis. See Holmer et al. (p. 41) (Holmer et al., 1996).

***Eoobolus acutulus*** Zhang, Zhang and Holmer sp. nov.

***Figure 2*** and ***Appendix 2*****—figures 5–7**, ***Appendix 3—table 2***.

**Etymology**. From the Latin '*acutulus*' (somewhat pointed), to indicate the slightly acuminate ventral valve with an acute apical angle. The gender is masculine.

**Holotype**. ELI-AJH S05 BT11 (***Appendix 2—figure 5E–H***), ventral valve.

**Paratype**. ELI-AJH S05 1-5-07 (***Appendix 2—figure 5M***), dorsal valve.

**Type locality**. Cambrian Series 2 Shuijingtuo Formation at the Aijiahe section (Zhang et al., 2016) near Aijiahe Village in Zigui County, north-western Hubei Province, China.

**Material**. 20 ventral and 3 dorsal valves, ranging from 715 μm to 1877 μm in length and from 562 μm to 1466 μm in width (***Appendix 2—figure 5*** and ***Appendix 3—table 2***) from Cambrian Series 2 Shuijingtuo Formation at the Aijiahe section, South China.

**Diagnosis**. Shell elongate oval with acuminate posterior margin, generally 128% longer than wide, biconvex. Metamorphic shell outlined by a pronounced halo, and ornamented with evenly distributed pits, while post-metamorphic shell ornamented with finely circular growth lines and large random pustules. Columnar shell structure. Ventral pseudointerarea orthocline with steep and narrow pedicle groove. Propareas small, with flexure lines. Posterolateral, umbonal and anterior muscle scars, and pedicle nerve weakly impressed. *Vascula lateralia* indefinite. Dorsal pseudointerarea orthocline. Propareas not raided. Median groove wide, poorly defined laterally, lacking flexure lines, short. Dorsal visceral area weakly impressed.

**Description**. Shell slightly ventribiconvex, elongate oval with acuminate posterior margin and rectimarginate anterior commissure, about 128% as long as wide (***Appendix 2—figure 5***). Metamorphic shell ornamented by regularly disposed hemispherical pits, uniform in size of about 0.6 μm, ranging from 0.2 μm to 1.1 μm in diameter (***Appendix 2—figure 6F–H*** and ***Appendix 3—table 2***). The pronounced halo marks the boundary between metamorphic shell and post-metamorphic shell (***Appendix 2—figure 6A*** and ***B***). Neanic shell indefinite. Post-metamorphic shell fully covered with finely and densely concentric growth lines superposed by elongate pustulose ornamentation, (***Appendix 2—figures 5C***, ***6I*** and ***J***). Elongate pustules have mean length of 11.8 μm, ranging from 8 μm to 21 μm, and mean width of 4.7 μm (***Appendix 3—table 2***), packed in concentric rows.

Ventral valve, acuminate with an acute apical angle of 83° on average ranging from 73° to 89°, elongate oval, and gently convex in sagittal profile, about length 130% width and depth 21% length, with maximum width slightly anterior to mid-length and maximum height slightly posterior to mid-length (***Appendix 3—table 2***). Pseudointerarea, orthocline, occupying 25% valve length and 69% valve width, with a deep, narrow, subtriangular pedicle groove of about 341 μm in length and 251 μm in width, with about 43% length of the pseudointerarea (***Appendix 2—figure 7A–C*** and ***G***). Lateral sides of the pedicle groove slightly divergent anteriorly at about 17°. Propareas narrow, raised above the valve floor, divided into two parts by robust flexure lines, while the internal part of proparea is wider than the external part (***Appendix 2—figure 6A*** and ***F***). Ventral valve interior with a vestigial visceral area, bisected by the divergent pedicle nerve impression terminated anteriorly at about 46% of valve length (***Appendix 2—figure 7C***). Paired posterolateral muscle scars weakly developed underneath the raised propareas (***Appendix 2—figure 7B***). Anterior and umbonal muscle scars weakly developed only on large valves (***Appendix 2—figure 7A*** and ***C***). *Vascula lateralia* indefinite. Metamorphic shell 229 μm long and 255 μm wide on average, bounded by a pronounced halo (***Appendix 2—figure 6A–C***). Possible protegulum noted as a distinct mound about 50 μm across, located posteromedially and bearing faint folds anterolaterally (***Appendix 2—figure 6C***).

Dorsal valve with apical angle of 90° on average, elongate oval, about 128% longer than wide, with maximum width slightly anterior to mid-length (electronic supplementary material, table S2). Pseudointerarea, wide, orthocline, occupying 19% valve length and 67% valve width, with a wide, shallow, subtriangular median groove of about 165 μm in length, with about 75% length of the pseudointerarea (***Appendix 2—figure 7H***). Median groove outline poorly diagnosed on pseudointerarea. Propareas very narrow. Dorsal valve interior with a vestigial visceral area. Median ridge vestigial (***Appendix 2—figure 5M***). Muscle systems poorly diagnosed.

Shell structure stratiform, consisting of primary laminated layer and secondary columnar layer (***Figure 2***). The primary laminated layer is composed of compact apatite, about 1.5 μm thick (**Figure 2*G***), while the secondary layer is composed of multi stacked sandwich columnar units (maximum 13 units), including numerous columns disposed orthogonally between a pair of stratiform lamellae (**Figure 2*B***, ***C***, ***J*** and ***K***; ***Appendix 2—figure 7J***). The hollow space in the columns and between stratiform lamellae of columnar units probably indicates the rich composition of organic material (***Figure 2F–H***). Columns are quite small about 2.4 μm in diameter, ranging from 1.2 μm to 3.2 μm, and about 4 μm in height, ranging from 1.7 μm to 6.5 μm. The central canal in the column small, ranging from 0.4 μm to 1 μm in diameter. The space between stratiform lamellae of stacked columnar units, thin, around 0.6 μm, while the stratiform lamellae about 1.2 μm in thickness (***Appendix 3—table 3***).
