## Appendix 2-figures and figure captions for "Evolution and diversity of biomineralized columnar architecture in early Cambrian phosphatic-shelled brachiopods"

**Supplementary figures and figure captions**

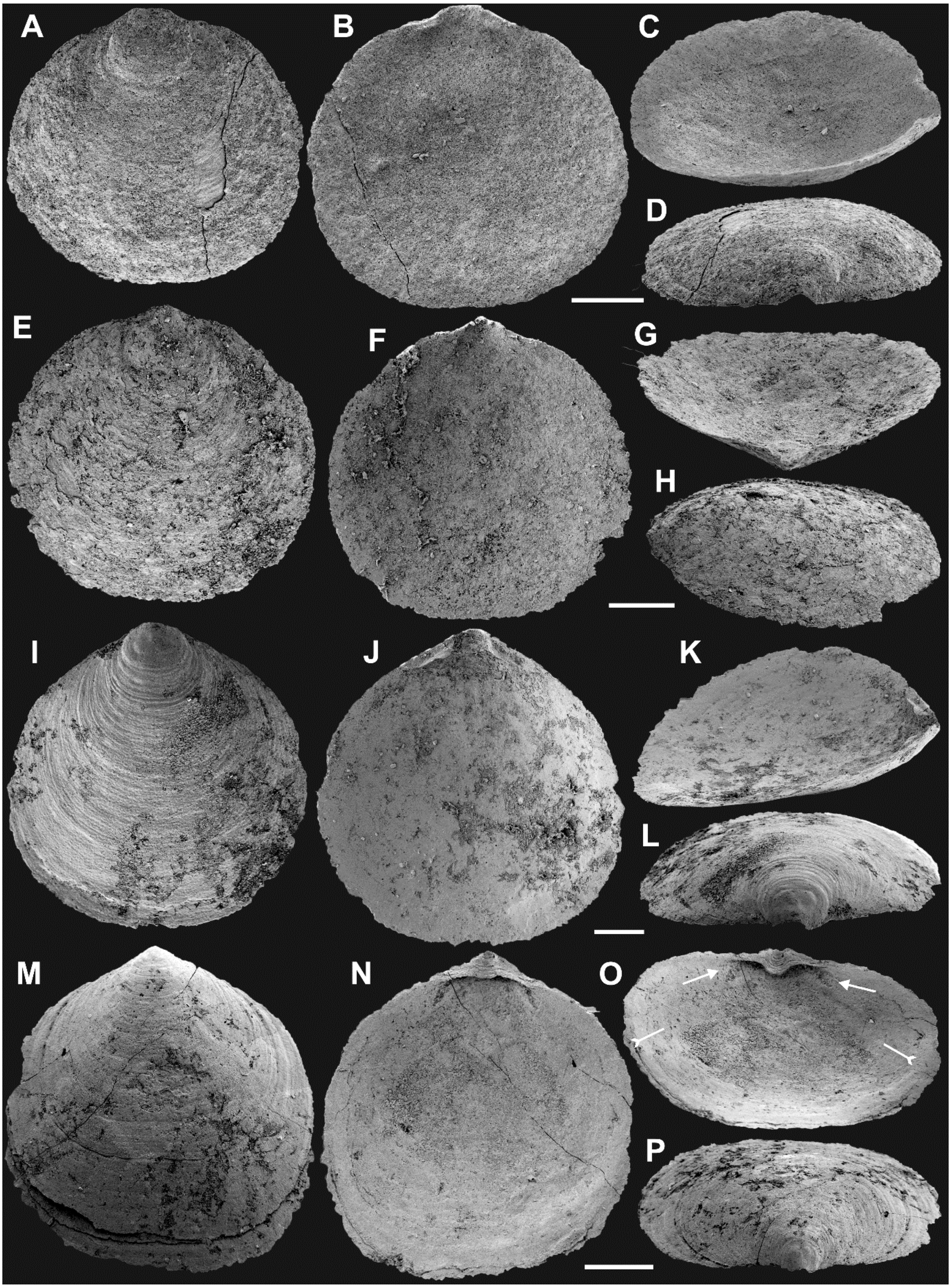


**Appendix 2—figure 1.** Ventral valves of *Latusobolus xiaoyangbaensis* gen. et sp. nov. from the Cambrian Series 2 Shuijingtuo Formation in southern Shaanxi, South China. **A-D**, juvenile with weakly developed pseudointerarea, ELI-XYB S5-1 BR01. **E-H**, juvenile with weakly developed pseudointerarea, ELI-XYB S5-1 BR02. **I-J**, large valve with slightly elevated pseudointerarea, ELI-XYB S5-1 BR04. **M-P**, mature valve with developed posterolateral muscle scars (arrows) and rudiment of *vascula lateralia* (tailed arrows), ELI-XYB S5-1 BR09. Scale bars: **A-L**, 200 μm; **M-P**, 500 μm.

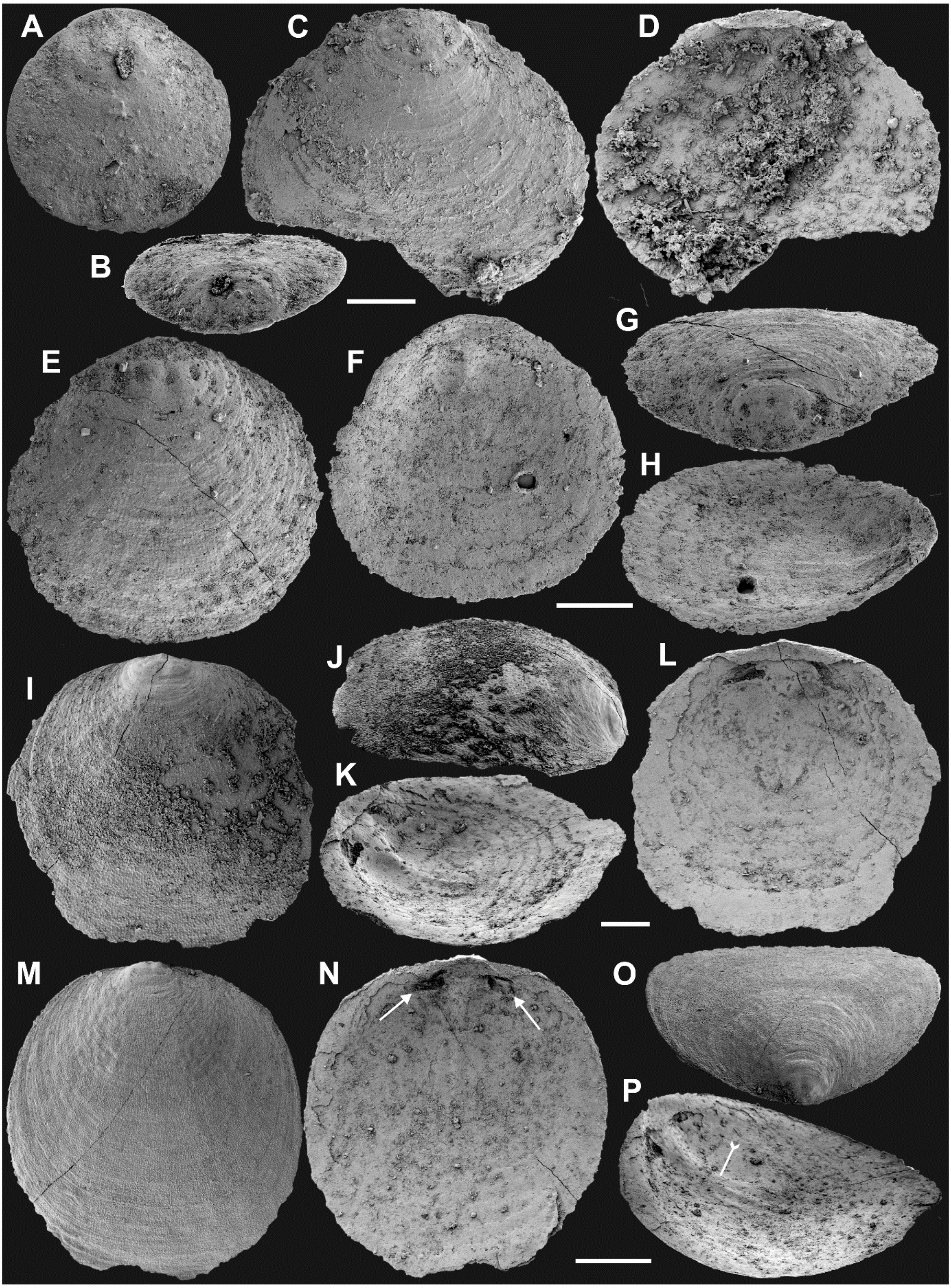


**Appendix 2—figure 2.** Dorsal valves of *Latusobolus xiaoyangbaensis* gen. et sp. nov. from the Cambrian Series 2 Shuijingtuo Formation in southern Shaanxi, South China. **A-B**, small juvenile with weakly developed pustular ornamentation, ELI-XYB S5-1 BS15. **C-D**, juvenile, ELI-XYB S4-2 BO08. **E-H**, juvenile with rudiment of median ridge, ELI-XYB S4-2 BS09. **I-L**, large valve, ELI-XYB S4-2 BO13. **M-P**, mature valve with weakly developed paired posterolateral muscle scars (arrows), median ridge and pair of submedian ridges bisecting dorsal visceral field (tailed arrow), ELI-XYB S4-2 BO11. Scale bars: **A-D**, 100 μm; **E-L**, 200 μm; **M-P**, 500 μm.

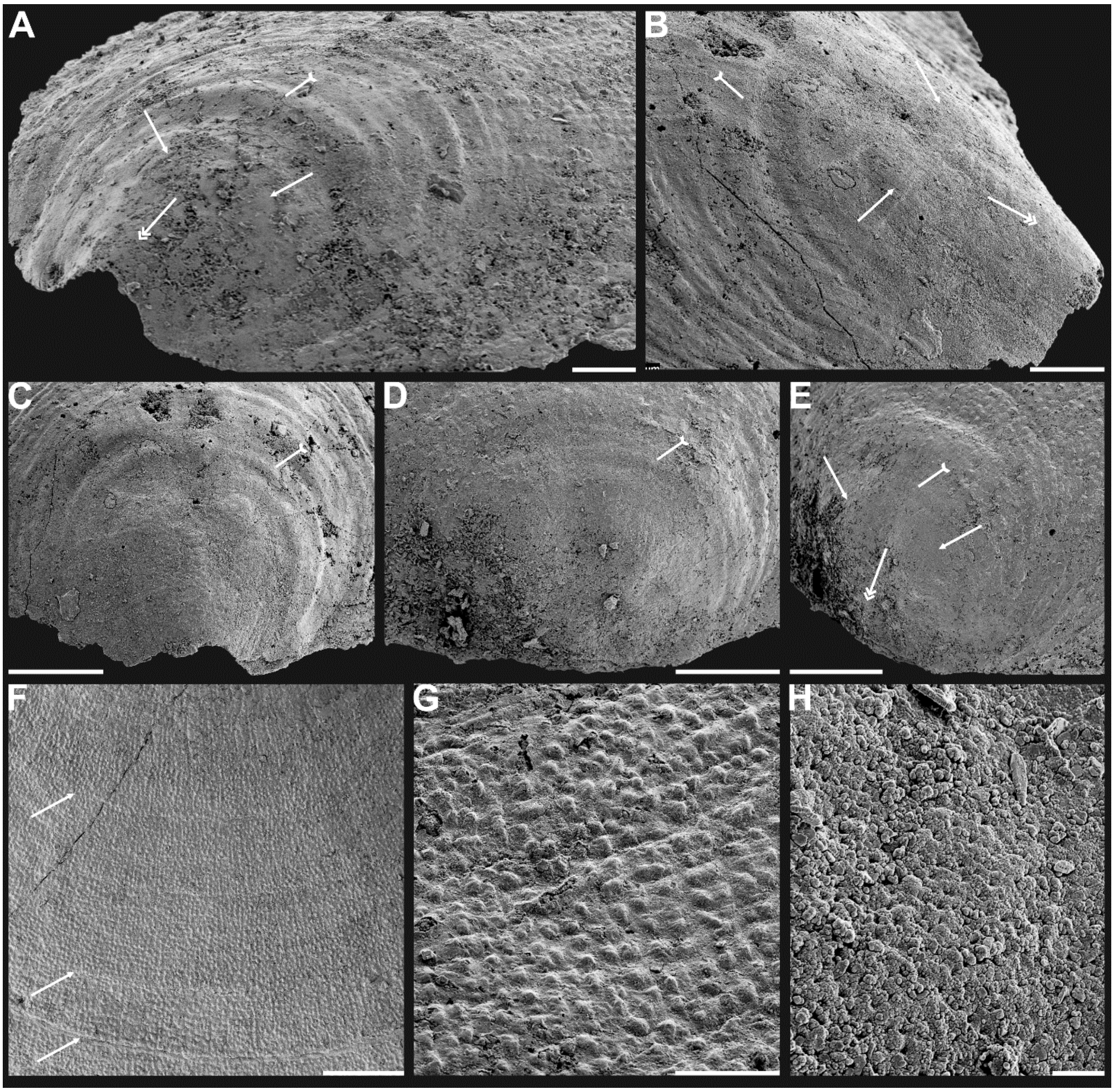


**Appendix 2—figure 3.** Shell characters and ornamentation of *Latusobolus xiaoyangbaensis* gen. et sp. nov. from the Cambrian Series 2 Shuijingtuo Formation in southern Shaanxi, South China. **A**, enlarged ventral metamorphic shell, noting the developed halo by tailed arrow, protegulum by double-headed arrow and brephic lobes by arrows, ELI-XYB S4-3 AU10. **B-C**, ventral valve, ELI-XYB S4-2 BO09. **B**, metamorphic shell of mature valve, noting the halo by tailed arrow, protegulum by double-headed arrow and brephic lobes by arrows. **C**, posterior view, noting the halo by tailed arrow. **D-G**, dorsal valve, ELI-XYB S4-2 BO11. **D**, metamorphic shell, noting the halo by tailed arrow. **E**, lateral dorsal view, noting the halo by tailed arrow, protegulum by double-headed arrow and brephic lobes by arrows. **F**, post-metamorphic pustules, note sparsely packed concentric growth lines by arrows. **G**, enlarged pustules. **H**, enlargement of metamorphic pitted ornament, ELI-XYB S4-2 BO08. Scale bars: **A**, **B**, **G**, 50 μm; **C-E**, 100 μm; **F**, 200 μm; **H**, 5 μm.

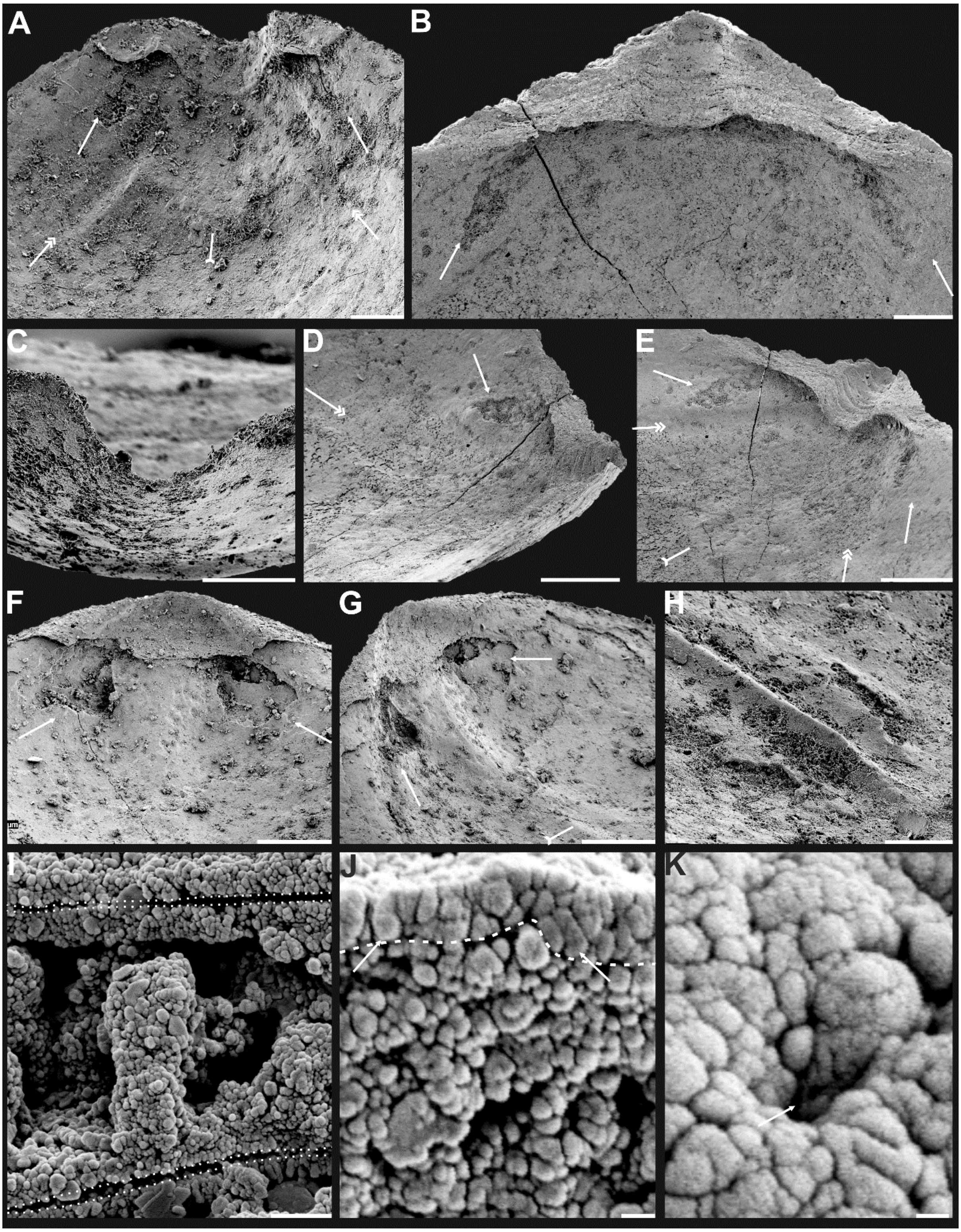


**Appendix 2—figure 4.** Internal characters and shell ultrastructures of *Latusobolus xiaoyangbaensis* gen. et sp. nov. from the Cambrian Series 2 Shuijingtuo Formation in southern Shaanxi, South China. **A-B**, ventral valve, ELI-XYB S4-3 AU10. A, interior showing posterolateral muscle scars below propareas by arrows, pedicle nerve by tailed arrow, *vascula lateralia* by double-headed arrows. **B**, posterior view of pedicle groove. **C-E**, ventral valve, ELI-XYB S5-1 BR09. **C**, interior view showing triangular pseudointerarea and paired posterolateral muscle scars by arrows. **D-E**, lateral view, note elevated pseudointerarea with posterolateral muscles underneath by arrows, pedicle nerve by tailed arrow, *vascula lateralia* by double-headed arrows. **F-G**, dorsal valve, noting paired umbonal muscle scars by arrows, median ridge by tailed arrow, ELI-XYB S4-2 BO11. **H**, enlargement of terminal of median ridge, ELI-XYB S5-1 BS07. **I**, ventral, enlarged columnar architecture, outlining organic

membranes between the stratiform lamellae of stacked sandwich columnar units, ELI-XYB S5-1 BS01. **J**, nanoscale apatite spherules of granule aggregations (arrows) of ventral shell, note primary-secondary layer boundary by dashed line, ELI-XYB S4-2 BO12. **K**, enlarged nanoscale spherules of pitted ornament (arrow) on primary layer, ELI-XYB S4-2 BO08. Scale bars: **A**, **B**, **H**, 50 μm; **C**, 100 μm; **D-G**, 200 μm; **I**, 2 μm; **J**, 500 nm; **K**, 200 nm.

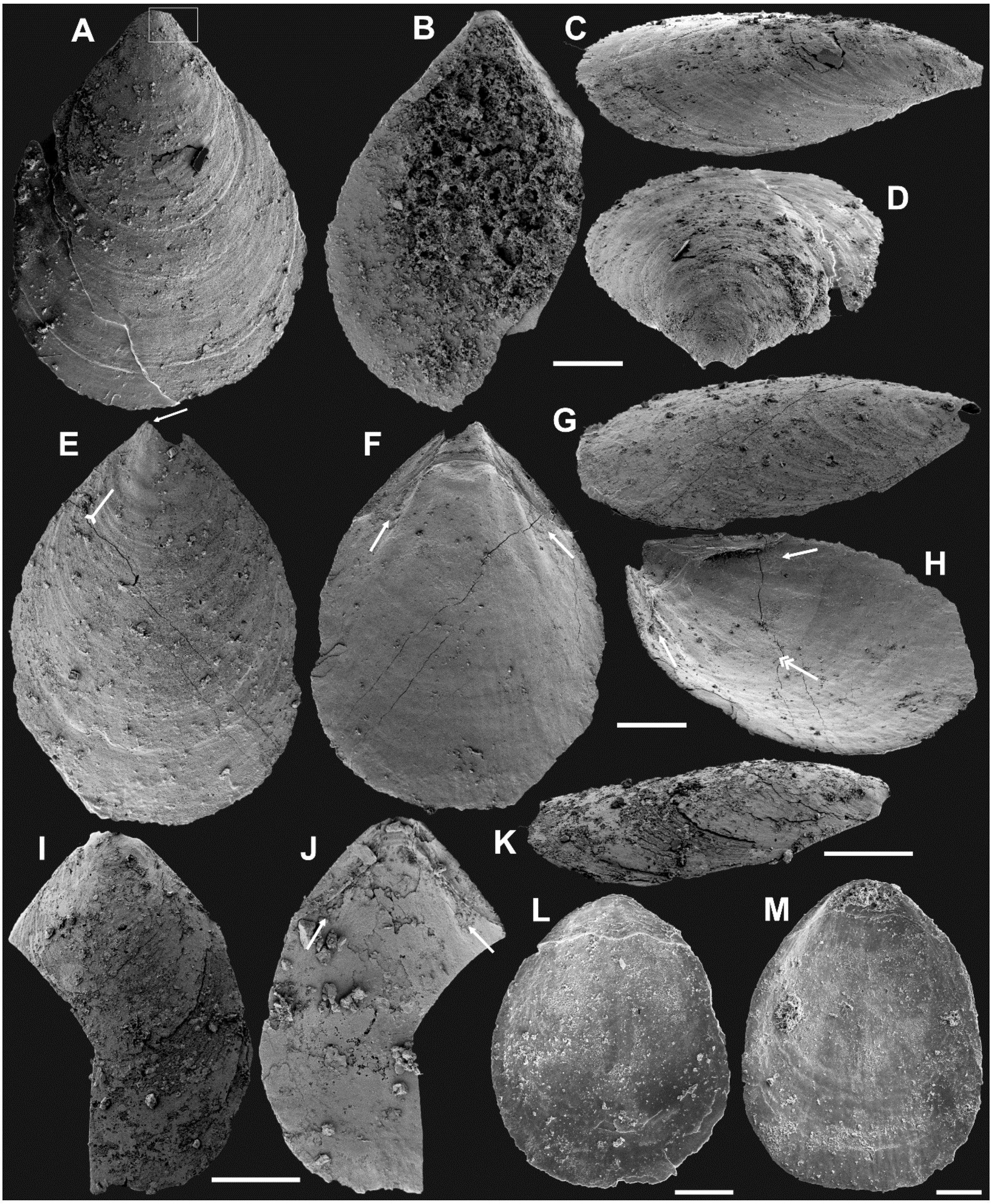


**Appendix 2—figure 5.** Ventral and dorsal valves of *Eoobolus acutulus* sp. nov. from the Cambrian Series 2 Shuijingtuo Formation in Three Gorges areas, South China. **A-D**, juvenile with weakly developed pseudointerarea, box indicates area in appendix 2—figure 6D, ELI-WJP 7 CE05. **E-H**, juvenile, note slightly developed pseudointerarea, the halo by tailed arrow, paired posterolateral muscle scars by arrows, pedicle nerve by double-headed arrow, ELI-AJH S05 BT11. **I-K**, mature valve, developed paired posterolateral muscle scars (arrows) underneath elevated pseudointerarea, ELI-AJH S05 BT14. **L-M**, dorsal valve with rudiment of median ridge, ELI-AJH 1-5-01, ELI-AJH 1-5-07. Scale bars: **A-H**, **L**, **M**, 200 μm; **I-K**, 400 μm.

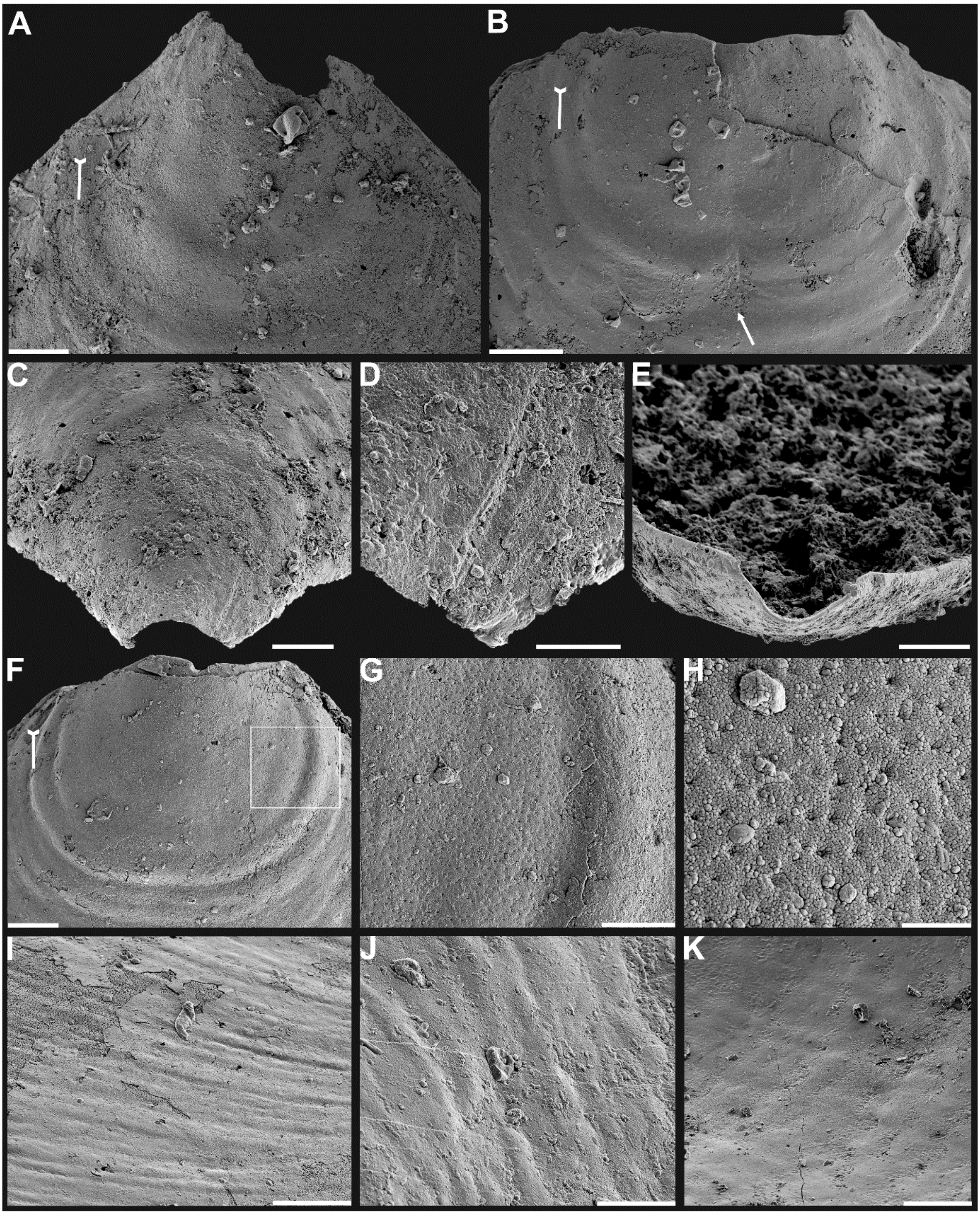


**Appendix 2—figure 6.** Shell characters and ornamentation of *Eoobolus acutulus* sp. nov. from the Cambrian Series 2 Shuijingtuo Formation in Three Gorges areas, South China. **A**, enlarged ventral metamorphic shell, noting the halo by tailed arrow, ELI-AJH S05 BT11. **B**, enlarged ventral metamorphic shell, noting the halo by tailed arrow and drape structures outside the halo by arrow, ELI-AJH S05 BT05. **C-E**, ventral valve, ELI-WJP 7 CE05. **C**, posterior ventral view of metamorphic shell. **D**, fine ridges on the margin of protegulum. **E**, posterior view of pedicle groove. **F-H**, ventral valve, ELI-AJH ST 8-2-3 BT04. **F**, metamorphic shell, noting the halo by tailed arrow, box indicates area in **G**. **G**, metamorphic pits. **H**, enlarged pits. **I**, dense concentric growth lines on ventral external surface, ELI-AJH 8-2-3 BT04. **J**, elongate pustular ornamentation on external surface, ELI-AJH 8-2-3 BT03. **K**, elongate pustule ornamentation on interior, ELI-AJH 8-2-3 BT04. Scale bars: **A-C**, **E**, **F**, **I**, **K**, 50 μm; **D**, **G**, 20 μm; **H**, 5 μm; **J**, 10 μm.

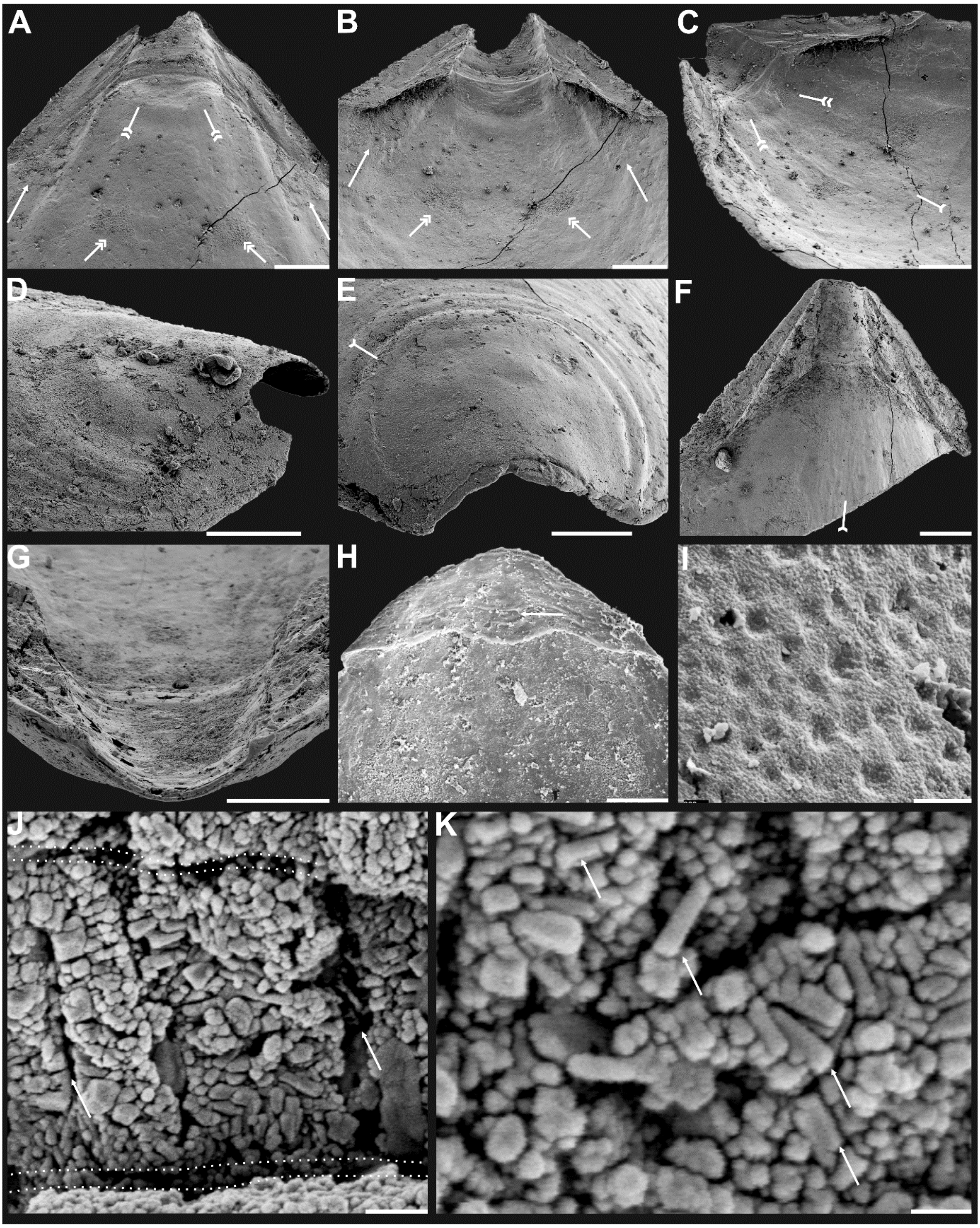


**Appendix 2—figure 7**. Shell characters and ultrastructures of *Eoobolus acutulus* sp. nov. from the Cambrian Series 2 Shuijingtuo Formation in Three Gorges areas, South China. **A-D**, ventral valve, ELI-AJH SJT S05 BT11. **A-C**, interior noting elevated pseudointerarea with posterolateral muscles underneath by arrows, pedicle nerve by tailed arrow, anterior muscle scars by double-headed arrows, umbonal muscle scars by double-tailed arrows. **D**, lateral view of metamorphic shell. **E**, posterior view of ventral metamorphic shell, note the halo by tailed arrow, ELI-AJH SJT S05 BT04. **F-G**, ventral valve, ELI-AJH SJT 8-2-3 BT02. **F**, pseudointerarea of a large valve, noting pedicle nerve by tailed arrow. **G**, posterior view of pedicle groove. **H**, dorsal pseudointerarea, ELI-AJH 1-5-01. **I**, enlarged pitted ornament on primary layer, ELI-AJH 8-2-2 Lin005. **J-K**, ventral valve, ELI-AJH S05 BT12. **J**, apatite spherules of granule aggregations, note organic canals of columns by arrows, outlining organic membranes between the stratiform lamellae of stacked sandwich column units. **K**, enlarged

nanoscale spherules of granule aggregations, note elongate rods by arrows. Scale bars: **A-C**, **G**, **H**, 100 µm; **D**, 50 µm; **E**, **F**, 200 µm; **I**, 2 µm; **J**, 1 µm; **K**, 400 nm.
