## Appendix 3-table 1 for "Evolution and diversity of biomineralized columnar architecture in early Cambrian phosphatic-shelled brachiopods"

**Appendix 3—table 1**. Average dimensions and ratios of ventral and dorsal valves of *Latusobolus xiaoyangbaensis* gen. et sp. nov. from the Cambrian Series 2 Shuijingtuo Formation, South China.

| V | L | W | H | $L_m$ | $L_{ms}$ | $W_{ms}$ | $L_n$ | $L_p$ | $W_p$ | $L_{pl}$ | $W_{pl}$ | $L_g$ | $W_g$ | $L_{p-i}$ | $W_{p-i}$ | $L_{p-e}$ | $W_{p-e}$ | A | $A_g$ | Pi | Pu | L/W | H/L |
|---|---|---|---|---|---|---|---|---|---|---|---|---|---|---|---|---|---|---|---|---|---|---|---|
| N | 17 | 15 | 20 | 11 | 19 | 19 | 5 | 14 | 13 | 6 | 5 | 17 | 20 | 12 | 11 | 12 | 11 | 19 | 14 | 28 | 40 | 14 | 17 |
| Mean | 1244 | 1237 | 239 | 655 | 223 | 275 | 523 | 165 | 615 | 321 | 634 | 112 | 120 | 157 | 29 | 277 | 51 | 129° | 34° | 0.5 | 6.5 | 95.17% | 18.02% |
| Min | 775 | 806 | 111 | 418 | 169 | 206 | 326 | 70 | 320 | 168 | 456 | 42 | 85 | 46 | 8 | 105 | 9 | 117° | 24° | 0.3 | 2.3 | 90.43% | 11.58% |
| Max | 2325 | 2417 | 519 | 1460 | 267 | 327 | 759 | 316 | 929 | 434 | 857 | 201 | 172 | 303 | 55 | 516 | 122 | 142° | 51° | 0.7 | 12.6 | 98.07% | 24.64% |
| Median | 983 | 998 | 198 | 520 | 223 | 275 | 427 | 152 | 602 | 329 | 558 | 99 | 120 | 137 | 29 | 255 | 45 | 126° | 32° | 0.5 | 6.4 | 95.99% | 17.09% |
| SD | 562 | 517 | 112 | 326 | 25 | 25 | 189 | 75 | 196 | 109 | 176 | 53 | 25 | 89 | 15 | 126 | 29 | 8 | 9 | 0.1 | 1.8 | 2.03% | 3.94% |

| V | $L_m$/L | $L_{ms}$/L | $L_{ms}$/$W_{ms}$ | $L_n$/L | $L_p$/L | $W_p$/W | $L_p$/$W_p$ | $L_g$/$L_p$ | $W_g$/$W_p$ | $L_g$/$W_g$ | $L_{pl}$/L | $W_{pl}$/W | $L_{p-i}$/$L_{p-e}$ | $W_{p-i}$/$W_{p-e}$ | $A_g$/A |
|---|---|---|---|---|---|---|---|---|---|---|---|---|---|---|---|
| N | 11 | 15 | 19 | 4 | 12 | 10 | 13 | 14 | 13 | 17 | 5 | 5 | 12 | 11 | 14 |
| Mean | 56.67% | 20.77% | 81.21% | 36.02% | 11.45% | 44.18% | 24.43% | 64.22% | 20.38% | 87.27% | 18.19% | 38.27% | 55.66% | 60.48% | 27.18% |
| Min | 53.50% | 8.60% | 68.81% | 32.65% | 7.79% | 35.75% | 18.09% | 42.41% | 15.68% | 49.41% | 13.77% | 28.00% | 20.63% | 30.51% | 18.44% |
| Max | 62.80% | 31.87% | 94.84% | 43.20% | 14.40% | 55.21% | 30.09% | 83.19% | 26.56% | 146.72% | 21.96% | 51.60% | 73.01% | 88.89% | 42.86% |
| Median | 56.68% | 22.84% | 80.77% | 34.13% | 11.31% | 43.01% | 26.14% | 62.16% | 19.96% | 79.84% | 17.89% | 39.21% | 56.86% | 57.69% | 24.07% |
| SD | 2.69% | 8.36% | 7.15% | 4.90% | 2.28% | 6.84% | 4.04% | 10.66% | 3.35% | 28.62% | 3.03% | 8.96% | 15.44% | 18.96% | 7.91% |

| D | L | W | H | $L_m$ | $L_{ms}$ | $W_{ms}$ | $L_r$ | $L_p$ | $W_p$ | $L_u$ | $W_u$ | $L_g$ | $W_g$ | A | $A_g$ | Pi | Pu | L/W | H/L | $L_m$/L | $L_{ms}$/L | $L_{ms}$/$W_{ms}$ |
|---|---|---|---|---|---|---|---|---|---|---|---|---|---|---|---|---|---|---|---|---|---|---|
| N | 16 | 15 | 16 | 12 | 10 | 10 | 9 | 11 | 11 | 5 | 5 | 7 | 5 | 13 | 4 | 17 | 41 | 15 | 16 | 12 | 10 | 10 |
| Mean | 1116 | 1240 | 229 | 569 | 198 | 260 | 746 | 112 | 579 | 283 | 734 | 104 | 292 | 136° | 112° | 0.5 | 6.3 | 92.6% | 19.5% | 55.6% | 18.6% | 76.0% |
| Min | 624 | 669 | 107 | 365 | 150 | 227 | 359 | 50 | 305 | 178 | 544 | 61 | 139 | 115° | 101° | 0.3 | 3.1 | 86.5% | 15.0% | 49.9% | 9.7% | 63.0% |
| Max | 2107 | 2190 | 550 | 1003 | 238 | 309 | 1243 | 205 | 1091 | 431 | 880 | 160 | 533 | 158° | 127° | 0.7 | 11.4 | 97.8% | 26.1% | 60.7% | 29.6% | 89.1% |
| Median | 897 | 1056 | 176 | 498 | 206 | 254 | 603 | 95 | 502 | 270 | 835 | 93 | 287 | 140° | 111° | 0.5 | 6.4 | 93.4% | 18.2% | 56.7% | 17.9% | 75.3% |
| SD | 498 | 526 | 140 | 198 | 33 | 29 | 357 | 48 | 265 | 104 | 172 | 41 | 158 | 13 | 11 | 0.1 | 1.8 | 3.7% | 3.5% | 3.1% | 7.0% | 9.6% |

| D | $L_r$/L | $L_p$/L | $W_p$/W | $L_g$/$L_p$ | $L_g$/$W_g$ | $W_g$/$W_p$ | $L_u$/L | $W_u$/W | $A_g$/A |
|---|---|---|---|---|---|---|---|---|---|
| N | 9 | 11 | 11 | 7 | 5 | 5 | 5 | 5 | 3 |
| Mean | 52.9% | 8.8% | 41.0% | 83.6% | 34.8% | 45.2% | 16.2% | 39.4% | 76.7% |
| Min | 32.4% | 6.3% | 35.6% | 78.0% | 27.5% | 30.9% | 12.8% | 36.9% | 72.2% |
| Max | 61.4% | 13.0% | 50.4% | 88.8% | 44.6% | 62.7% | 22.3% | 42.1% | 82.5% |
| Median | 53.6% | 8.5% | 39.9% | 84.9% | 36.2% | 45.0% | 16.2% | 39.3% | 75.4% |
| SD | 9.3% | 1.9% | 5.0% | 4.1% | 7.0% | 11.4% | 3.9% | 1.9% | 5.3% |

All measurements are in µm. Abbreviations: A, apical angle; $A_g$, angle of ventral pedicle groove or dorsal median groove; D, dorsal valve; Pi, diameter of pitted structures; Pu, diameter of pustules; V, ventral valve. L, length; W, width; H, height of valve where not specified, and of elements: g, ventral pedicle groove or dorsal median groove; m, valve length at the maximum width; r, median ridge; ms, metamorphic shell; p, pseudointerarea; p-i, inner part of proparea; p-o, outer part of proparea; pl, posterolateral muscle scars; pu, pustules, n, pedicle nerve; u, umbonal muscle scars.
