## Appendix 3-table 2 for "Evolution and diversity of biomineralized columnar architecture in early Cambrian phosphatic-shelled brachiopods"

**Appendix 3—table 2**. Average dimensions and ratios of ventral and dorsal valves of *Eoobolus acutulus* sp. nov. from the Cambrian Series 2 Shuijingtuo Formation, South China.

| V | L | W | H | $L_m$ | $L_{ms}$ | $W_{ms}$ | $L_n$ | $L_p$ | $W_p$ | $L_{pl}$ | $W_{pl}$ | $L_g$ | $W_g$ | $L_{p-i}$ | $W_{p-i}$ | $L_{p-e}$ | $W_{p-e}$ | A | $A_g$ | Pi | $L_{pu}$ | $W_{pu}$ | L/W |
|---|---|---|---|---|---|---|---|---|---|---|---|---|---|---|---|---|---|---|---|---|---|---|---|
| N | 13 | 10 | 5 | 11 | 11 | 12 | 1 | 9 | 7 | 2 | 2 | 10 | 12 | 11 | 12 | 10 | 11 | 20 | 11 | 68 | 18 | 18 | 10 |
| Mean | 1211 | 979 | 285 | 756 | 229 | 255 | 539 | 594 | 1079 | 459 | 673 | 341 | 251 | 443 | 155 | 1066 | 137 | 83° | 17° | 0.6 | 11.8 | 4.7 | 129.53% |
| Min | 715 | 562 | 178 | 442 | 181 | 191 | 539 | 178 | 509 | 329 | 548 | 102 | 141 | 128 | 30 | 348 | 43 | 73° | 6° | 0.2 | 8.0 | 2.7 | 122.90% |
| Max | 1877 | 1466 | 432 | 1110 | 277 | 301 | 539 | 1096 | 1877 | 589 | 798 | 629 | 567 | 1027 | 406 | 2933 | 312 | 89° | 31° | 1.1 | 21.0 | 7.2 | 136.16% |
| Median | 1170 | 992 | 295 | 784 | 237 | 263 | 539 | 487 | 974 | 459 | 673 | 366 | 211 | 382 | 150 | 908 | 135 | 84° | 17° | 0.6 | 10.9 | 4.8 | 130.36% |
| SD | 335 | 280 | 96 | 210 | 37 | 30 | 0 | 328 | 463 | 184 | 177 | 187 | 117 | 275 | 105 | 763 | 78 | 5 | 9 | 0.2 | 3.4 | 1.3 | 4.80% |

| V | H/L | $L_m$/L | $L_{ms}$/L | $L_{ms}$/$W_{ms}$ | $L_n$/L | $L_p$/L | $W_p$/W | $L_p$/$W_p$ | $L_g$/$L_p$ | $W_g$/$W_p$ | $L_g$/$W_g$ | $L_{pl}$/L | $W_{pl}$/W | $L_{p-i}$/$L_{p-e}$ | $W_{p-i}$/$W_{p-e}$ | $A_g$/A |
|---|---|---|---|---|---|---|---|---|---|---|---|---|---|---|---|---|
| N | 4 | 11 | 8 | 10 | 1 | 4 | 3 | 6 | 5 | 6 | 8 | 2 | 2 | 9 | 10 | 10 |
| Mean | 21.49% | 60.79% | 19.53% | 92.06% | 46.03% | 24.73% | 68.79% | 53.52% | 43.40% | 20.12% | 155.51% | 31.40% | 62.94% | 40.92% | 103.65% | 21.54% |
| Min | 16.05% | 55.85% | 10.07% | 73.83% | 46.03% | 19.16% | 56.56% | 47.59% | 38.81% | 10.28% | 69.01% | 28.10% | 62.15% | 32.81% | 69.77% | 6.74% |
| Max | 25.46% | 67.01% | 25.31% | 109.91% | 46.03% | 28.70% | 75.86% | 59.94% | 57.30% | 31.04% | 229.02% | 34.71% | 63.72% | 56.03% | 156.21% | 35.62% |
| Median | 22.23% | 61.34% | 20.51% | 95.51% | 46.03% | 25.53% | 73.95% | 53.68% | 40.33% | 19.02% | 155.29% | 31.40% | 62.94% | 39.80% | 93.02% | 21.94% |
| SD | 4.62% | 3.01% | 4.87% | 11.80% | #DIV/0! | 4.02% | 10.64% | 5.66% | 7.87% | 7.99% | 57.93% | 4.68% | 1.11% | 7.54% | 30.35% | 10.87% |

| D | L | W | $L_m$ | $L_p$ | $W_p$ | $L_g$ | A | L/W | $L_m$/L | $L_p$/L | $W_p$/W | $L_g$/$L_p$ |
|---|---|---|---|---|---|---|---|---|---|---|---|---|
| N | 2 | 2 | 2 | 2 | 2 | 2 | 2 | 2 | 2 | 2 | 2 | 2 |
| Mean | 1203 | 939 | 723 | 229 | 637 | 165 | 90° | 127.7% | 59.7% | 18.8% | 67.4% | 74.7% |
| Min | 1015 | 817 | 578 | 178 | 519 | 156 | 87° | 124.2% | 56.9% | 17.5% | 63.5% | 61.8% |
| Max | 1390 | 1060 | 867 | 280 | 755 | 173 | 93° | 131.1% | 62.4% | 20.1% | 71.2% | 87.6% |
| Median | 1203 | 939 | 723 | 229 | 637 | 165 | 90° | 127.7% | 59.7% | 18.8% | 67.4% | 74.7% |
| SD | 265 | 172 | 204 | 72 | 167 | 12 | 4 | 4.9% | 3.8% | 1.8% | 5.4% | 18.3% |

All measurements are in µm. Abbreviations: A, apical angle; $A_g$, angle of ventral pedicle groove or dorsal median groove; D, dorsal valve; Pi, diameter of pitted structures; V, ventral valve. L, length; W, width; H, height of valve where not specified, and of elements: g, ventral pedicle groove or dorsal median groove; m, valve length at the maximum width; r, median ridge; ms, metamorphic shell; p, pseudointerarea; p-i, inner part of proparea; p-o, outer part of proparea; pl, posterolateral muscle scars; pu, pustules, n, pedicle nerve; u, umbonal muscle scars.
